# Cell-to-cell mobility of the stem cell inducing WUSCHEL transcription factor is controlled by a balance of transport and retention

**DOI:** 10.1101/2024.10.17.618816

**Authors:** Michael Fuchs, Thomas Stiehl, Anna Marciniak-Czochra, Jan U. Lohmann

## Abstract

Non-cell autonomous induction of stem cell fate is a shared feature across multicellular organisms, however the underlying mechanisms diverge substantially between the kingdoms of live. In plants, cell to cell mobility of transcription factors has emerged as a key paradigm. For the shoot apical meristem of the reference plant *Arabidopsis thaliana* it has been described that the translocation of the WUSCHEL homeodomain transcription factor from niche cells to stem cells is essential for their maintenance. Here we systematically investigate the function of diverse WUS alleles and leverage multispectral live cell imaging coupled to computational analysis and mechanistic mathematical modelling to show that WUSCHEL protein mobility is the result of balance between active transport and retention in niche cells and likely independent of the stem cell signal CLAVATA3. Importantly, we show that diffusion across cell layers of the meristem is not symmetrical, suggesting that there is unexpected complexity in cellular connections.

## Introduction

Post-embryonic development allows plants to show remarkable degrees of phenotypic plasticity in adaptation to constantly changing environmental conditions, providing the means to compensate for many disadvantages brought about by a sessile life-style. To this end, plants maintain pluripotent stem cells in specialized niche-tissues, called meristems, where they are subject to strict regulation: Cell proliferation and cell differentiation are balanced and, as a result, stem cell homeostasis is ensured throughout the plants entire life-cycle. The shoot apical meristem (SAM), which is situated at the top of the shoot, harbors stem cells that directly or indirectly contribute to the formation of almost all arial tissues. In *Arabidopsis thaliana*, their activity is, to a large extend, dependent on the non-cell autonomous activity of the homeodomain transcription factor WUSCHEL (WUS).

*WUS* mRNA is expressed in a sub-apical domain in the deeper cell layers of the meristem, called L3, and defines the organizing center (OC) ^1^. From the OC, WUS protein moves apically into the stem cells in the central zone (CZ) of the sub-epidermal (L2) and epidermal (L1) cell layers via cytoplasmic bridges called plasmodesmata ^2,3^. WUS acts directly in stem cells where it affects plant hormone signaling, including auxin and cytokinin, as well as expression of the stem cell marker *CLAVATA 3* (*CLV3*) ^4–9^. CLV3, a small peptide, in turn, is secreted from the stem cells and moves through the apoplast to the OC, where it initiates a signaling cascade to inhibit *WUS* expression and with that, indirectly, also its own expression ^9–14^. As a result, *WUS* and *CLV3* form a spatially separated negative feedback loop along the apical-basal axis in the center of the meristem that controls cell proliferation of the stem cells via constant balancing of *WUS* expression levels to maintain a small pool of pluripotent stem cells at the meristem tip. Recently, this model has been extended by the discovery of another negative feedback loop involving *CLAVATA3 / ESR-RELATED 40* (*CLE40*), a peptide closely related to *CLV3* and previously known for its role in regulating cell proliferation in the root meristem via restricting the expression of *WUSCHEL-RELATED HOMEOBOX 5* (*WOX5*) ^15–18^. In the shoot meristem, *CLE40* is expressed in the peripheral zone (PZ), complementing both the *CLV3* and *WUS* expression domains. Unlike CLV3, CLE40 itself is proposed to be an autocrine signaling molecule that acts via the activation of a currently unknown non-cell autonomous downstream factor to promote *WUS* expression in the OC ^19^. WUS protein, which is present within the cells of the OC as well as the stem cells in the CZ, in turn, represses *CLE40*, restricting its expression to the PZ ^19^. As a result, *CLE40* signaling conveys information on the size of the peripheral zone and the requirement for new cells to be integrated into organ primordia. Taken together, the interdependent activity of the antagonistic *WUS-CLV3* and *WUS-CLE40* feedback loops defines the size of the stem cell domain in all three dimensions: vertically, along the apical-basal axis (*WUS-CLV3*) and horizontally, along the center-periphery axis (*WUS-CLE40*).

Besides being a core component of the feedback loops that provide robustness to the shoot stem cell system, our growing understanding of stem cell regulatory networks has made it clear that *WUS* serves as a central integrator of local and global information on the physiological and developmental status of the plant, as well as of external environmental cues, both biotic and abiotic. One prime example is the response of the meristem to nitrogen, a key nutrient limiting plant growth under most natural settings. Upon nitrate uptake by the root, cytokinin signaling is activated in the shoot, which promotes WUS expression and in turn shoot meristem size and organ production rate ^20^.

Many activities of *WUS* are intrinsically tied to its ability to move from cell to cell in a symplastic manner: Preventing WUS movement from the OC to the CZ by blocking plasmodesmata in *CLV3* positive cells leads to stem cell depletion and meristem termination, similar to stem cell specific degradation of WUS protein ^2,7^. Conversely, bypassing WUS movement via miss-expression directly from the *CLV3* promoter triggers autocatalytic feedback expression of WUS, which ultimately leads to massive over-proliferation of the SAM ^8^. While the importance of protein mobility for *WUS* function has been clearly established and even though previous studies have suggested potential regulatory mechanisms, including homo-dimerization of WUS protein ^2,5,21^, nuclear-cytoplasmic partitioning ^21–23^, protein destabilization ^21–23^ and protein-protein interactions ^24–26^, we still lack a clear understanding of the quantitative regulation of WUS mobility and, most importantly, on how this process may be controlled in coordination with *WUS* transcriptional regulation.

Analysis of the movement capacities of WUS within the SAM has relied heavily on live-cell imaging of protein fusions with fluorescent tags. While this approach has led to the description of remarkable phenotypes, including drastic changes in meristem architecture as a result of altered WUS distribution patterns ^2^, it has been limited to mostly qualitative analysis of protein distribution by visual inspection or simple image quantifications. Importantly, in addition to the lack of reliable quantifications, the influence of various forms of WUS tagging has not been studied. Therefore, it seems reasonable to assume that some of the effects reported might be specific to a particular WUS allele and that more subtle variations in protein distribution -likely to be biologically significant in a system as highly regulated as the *WUS*-mediated maintenance of stem cells - may have escaped notice.

To address this problem, we have applied an integrated approach building on quantitative live cell imaging, unbiased comparison of tagged WUS alleles and mechanistic mathematical modeling. Leveraging these tools, we report here that tagging substantially affects WUS mobility and present evidence for nondirectional active transport of WUS, which is counteracted by protein retention in the L3. In line with previous results ^2,21^, our data suggests that the WUS homeodomain is necessary for active transport and that unstructured regions at the N- or C-terminus have important regulatory functions.

## Results

Fluorescent tags have become one of the primary methods for *in-vivo* analysis of protein behavior, localization and function. While the general usefulness of fluorescent tagging cannot be overstated, it is important to keep in mind that the attachment of a fluorophore may result in inadvertent side effects due to the size of the tag, sterical hinderance of functional domains or even changes in overall protein structure. In case of WUS, various tagged alleles have been used to study the *in-vivo* behavior of the protein and conflicting conclusions have been derived from a mostly visual analysis of individual transgenic lines ^2,21–23^. However, to date no systematic comparison of tagged WUS alleles with regards to functionality or protein distribution within the meristem has been performed. To address this deficiency, we created N-and C-terminal WUS protein fusions with green fluorescent protein (GFP), with and without the addition of a flexible serine-glycine linker (GFP-WUS, WUS-GFP, GFP-linker-WUS, WUS-linker-GFP) (Fig 1B), employing our GreenGate cloning system, which causes only minimal cloning scars ^27^, and used the resulting transgenes to complement *wus* null mutants. Plants homozygous for *wus* are unable to maintain a stable apical stem cell system, resulting in a stop-and-go mode of growth and development due to repetitive initiation and loss of stem cells, and in addition lack a functional set of reproductive organs, rendering them infertile ^28^. We established homozygous single-insertion rescue lines for all four fusion proteins and analyzed rescue efficiency, as well as protein distribution using live-cell imaging.

**Figure 1.**
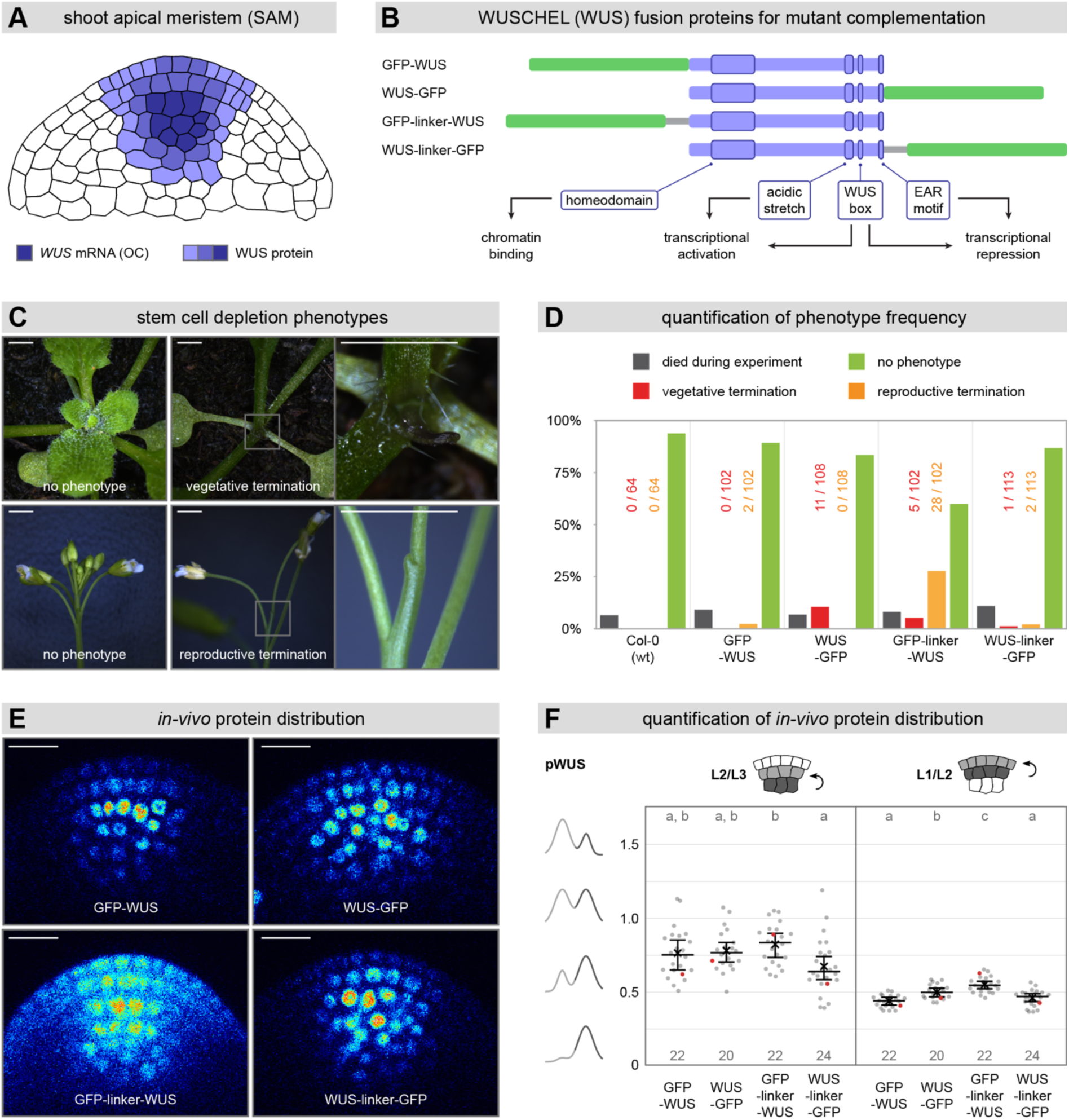
Systematic comparison of GFP tagged WUS alleles. **(A)** Schematic representation of *WUS* mRNA and WUS protein distribution in the shoot apical meristem (SAM). **(B)** Schematic comparison of different WUS fusion proteins, including the relative position of important N- and C-terminal protein domains. **(C)** Stem cell depletion phenotypes observed in *wus*-mutant complementation lines (T3). Scale bars represent a length of 2 mm. **(D)** Quantitative analysis of the rescue efficiency of different *wus*-mutant complementation lines (T3) compared to Col-0 wildtype (wt) plants. **(E)** Live-cell imaging of different *wus*-mutant complementation lines (T3). Images have been acquired from a native side-view perspective. All meristems shown here have been imaged with identical microscope settings. Scale bars represent a length of 15 µm. **(F)** Quantitative analysis of upwards mobility within different *wus*-complementation lines (T3) using ITQT. Red data points correspond to the meristems displayed in (E). Lines represent population median values with a 95% confidence interval. Crosses represent population mean values. Sample numbers are indicated below each population. Letters represent the result of pairwise comparisons by the ANOVA-TukeyHSD test (groups sharing a common letter are not significantly different at the 0.01 level).

While all WUS fusions were able to complement the *wus*-mutant phenotype to some extent and rescue lines were generally similar to wild-type plants, there were notable differences in the frequency of meristematic phenotypes. Importantly, these included stem cell depletion and termination of the meristem (Fig 1C), either at the vegetative stage before outgrowth of the main inflorescence shoot, or at the reproductive stage after bolting. Plants affected in vegetative development were usually able to produce a small number of true leaves before stem cells were depleted and development arrested. Formation of meristematic tissue was re-initiated subsequently and led to the disorganized outgrowth of atypical structures. Plants affected in reproductive development grew normal during vegetative development, but after bolting and outgrowth of the main inflorescence produced only a small number of floral organs before the meristem terminated. We observed no re-initiation of stem cells resulting in the formation of either additional flowers or atypical organ structures following meristem termination, however, the flowers produced before stem cell depletion appeared phenotypically normal and developed siliques.

These phenotypes, which we could not observe in Col-0 wild-type control plants (0/64), occurred at a low frequency in the GFP-WUS line (1.96% reproductive termination (2/102)) and the WUS-linker-GFP line (0.88% vegetative termination (1/113), 1.76% reproductive termination (2/113)) (Fig 1D). In contrast, the WUS-GFP line, showed a much higher fraction of plants with terminated vegetative meristems (10.19% vegetative termination (11/108)), while the GFP-linker-WUS line was affected in both vegetative and reproductive development (4.9% vegetative termination (5/102), 27.45% reproductive termination (28/102)). Taken together, these results suggested that different tagging strategies have a considerable influence on WUS function *in-vivo*, with GFP-WUS and WUS-linker-GFP exhibiting the highest frequency of rescued plants in our growth conditions.

Next, we wanted to test for correlation between this functional read-out and *in-vivo* protein distribution. To this end, we subjected our *wus*-rescue lines to confocal live-cell imaging, which was performed from a side-view perspective to maximize image resolution along the apical-basal axis of the meristem ^29^. In addition, we carefully adjusted microscopy settings for each sample to avoid over-saturation of fluorescence signal, allowing for subsequent quantitative analysis. Initial visual inspection of fluorescence images (Fig 1E) showed little differences between *wus*-rescue lines containing WUS tagged at the N-or C-terminus without a linker (GFP-WUS, WUS-GFP) and the line containing a C-terminal fusion with a linker (WUS-linker-GFP): All protein variants were present in the organizing center (OC), where the WUS promoter is active ^1^, as well as in the sub-epidermal (L2) and epidermal (L1) cell layers, where WUS protein moves to via plasmodesmata ^2,3^. The amount of protein gradually decreased from the L3 to the L1 and sub-cellular localization of the fusion proteins was predominantly nuclear, in line with previous reports ^2,3^. In contrast, the N-terminal WUS fusion protein with the same flexible serine-glycine linker (GFP-linker-WUS) appeared less nuclear and the gradient of protein from the L3 to the L1 was less steep, leading to an increased relative amount of protein in L2 and L1 in comparison to the other fusions (Fig 1E).

Together with the results on *in-vivo* functionality, these data suggested that fairly minor quantitative differences in WUS distribution may result in phenotypic variations, pointing to the need for a reliable method for precise and large-scale image quantification. We therefore developed a computational ‘Image Transformation and Quantification Tool’ (ITQT), which facilitates fluorescent signal quantification in a complex three-dimensional tissue by a sequence of semi-automated dimension reduction steps (Fig S1). ITQT is able to compensate for the curvature of the shoot meristem and to generate layer-specific fluorescence intensity readings within a cylindrical column around the OC along the apical-basal axis of the SAM. To this end, ITQT subjects every image to a polar transformation based on three circle-points, which results in the alignment of cells that belong to the same cell layer and prevents false allocation of fluorescent signal (Fig S1A). The aligned image stack (3D) is projected into a single plane (2D), before compressing the information into a single dimension (1D) (Fig S1B). The resulting pixel intensity curve is then used to visually identify the borders between cell layers, which are subsequently used to extract layer specific signal intensity from the two-dimensional image projection. In order to compare individual plants with the same genetic background as well as different lines – regardless of expression strength and laser-power applied while imaging – we calculated the intensity ratios between adjacent cell layers (L2/L3 and L1/L2). For this, we divided the fluorescence intensity of the more apical cell layer by the fluorescence intensity within the more basal layer: A low ratio (<1) indicated that the amount of protein in the apical layer was lower than in the basal layer, while a high ratio (>1) indicated the reverse. Relative differences in layer ratios (e.g., L2/L3) between different lines were interpreted as a measure for the differential ability of the respective fusion protein to move from the basal layer (e.g., L3) to the adjacent apical layer (e.g., L2).

Our quantitative analysis (Fig 1F) confirmed the visual impression of a decreasing L3-to-L1 gradient for all rescue lines. Interestingly however, even though all constructs were established as homozygous single-insertion lines, the variability within populations was generally high -but comparable between different lines - and a small number of plants showed a L2/L3 ratio >1, indicating that in these rare individuals, the amount of WUS fusion protein in the L2 was even higher than in the L3 where the *WUS* promoter is active. The median L2/L3 ratio was lowest for WUS-linker-GFP (L2/L3 median ratio 0.634, n = 24), followed by GFP-WUS (L2/L3 median ratio 0.746, n = 22), WUS-GFP (L2/L3 median ratio 0.760, n = 20) and GFP-linker-WUS (L2/L3 median ratio 0.829, n = 22) with the highest ratio. Statistical analysis revealed no significant differences between rescue lines, except for the direct comparison of WUS-linker-GFP and GFP-linker-WUS. Looking at the L1/L2 ratios, we observed the lowest median ratio for GFP-WUS (L1/L2 median ratio 0.434, n = 22), followed by WUS-linker-GFP (L1/L2 median ratio 0.464, n = 24), which were not significantly different from each other, as well as WUS-GFP (L1/L2 median ratio 0.493, n = 20) and GFP-linker-WUS (L1/L2 median ratio 0.539, n = 22). Variability was lower compared to the L2/L3 ratio and except for the direct comparison of GFP-WUS and WUS-linker-GFP, where we could see no statistically significant difference, all lines were significantly different from each other. Overall, our quantitative analysis suggests little difference in protein distribution between N-terminally tagged WUS protein without a linker (GFP-WUS) and C-terminally tagged WUS protein with a linker (WUS-linker-GFP). Both lines did not show any statistically significant difference, regardless of which layer ratio we analyzed. In contrast, using a C-terminal tag with no linker (WUS-GFP) slightly increased the amount of protein in L2 and L1, while the addition of a linker between an N-terminal tag and WUS (GFP-linker-WUS) drastically increased the amount of fusion protein in upper meristematic cell layers. Taken together, our quantitative phenotypic and microscopic analysis based on transgenes generated using the same vector system demonstrated that GFP-WUS and WUS-linker-GFP are equally suited for the analysis of WUS protein mobility. Incidentally, these alleles represent the two forms of WUS fusions most commonly used within the community. Having established ITQT as a highly sensitive and reproducible pipeline for large-scale image quantification in the SAM, we were curious to re-visit a number of questions central to understanding the mechanism behind the movement of any mobile protein: Is the mobility due to passive diffusion or active transport and to what extend is movement dependent on protein structure versus a specific cellular environment? To investigate differential mobility between meristematic cell layers, we expressed several fusion proteins from the *WUS* promoter, in order to assess basal to apical movement, from the OC to the CZ, as well as from the epidermal *MERISTEM LAYER 1* (*ML1*) promoter ^30^ to assess movement in the reverse direction, from the L1 to the OC. Based on our results described above, we decided to use C-terminal fusions containing a flexible serine-glycine linker and to analyze T1 populations rather than homozygous single-insertion lines, given the high variability observed in established lines. We reasoned that this approach would help to average possible effects of the genomic-insertion site on transgene expression and would prevent potential bias in selecting individuals for line establishment, consequently allowing to cover and display the full extent of natural variability. In addition, all plants were imaged from a top-view perspective, which compared to side-view imaging has a reduced imaging resolution along the apical-basal axis, but due to a simpler dissection and imaging procedure ^29^, allowed for the processing and analysis of a larger number of plants. Importantly, we also controlled for within-construct variation and between-construct variation by testing an alternative fluorophore (mNeonGreen instead of GFP), plant resistance cassette (Hygromycin instead of BASTA) and GreenGate destination vector backbone (pGGZ001 instead of pGGZ003). Since our analysis did not reveal any systematic differences in the behavior of WUS in these settings (Fig S2), we combined results obtained from different constructs with the same protein of interest (e.g., WUSΔbox-linker-GFP + WUSΔbox-linker-mNeonGreen = WUSΔbox-linker-FP) to further increase sample size and therefore improve statistical power.

In order to avoid phenotypic effects characteristic for WUS miss-expression, such as ectopic initiation of stem cells and over-proliferation, which both would result in drastic changes in overall meristem architecture, we made use of a WUS variant, termed WUSΔbox, that has been transcriptionally inactivated via mutation of its WUS box ^31^. This mutation of the WUS box has previously been reported to have no qualitative effect on WUS mobility ^2^, making Δbox variants powerful tools for the analysis of protein movement by allowing to induce alterations in protein distribution with minimal impact on endogenous WUS feedback regulation. We first analyzed the vertical distribution of WUSΔbox(-linker-FP) compared to 2xGFP, representing a similarly sized but freely diffusible control. 2xGFP tagged with a strong nuclear localization signal (2xGFP-NLS), which in a previous study has been reported to not move beyond its expression domain ^2^, was considered as a second, immobile control. Since a previous publication has suggested that sub-cellular partitioning between the nucleus and the cytoplasm is involved in controlling WUS distribution ^21^, we also characterized the distribution of WUSΔbox tagged with a small nuclear export signal (NES) at its N-terminus (NES-WUSΔbox(-linker-FP)). In addition, we analyzed the mobility of MiniMeΔbox(-linker-FP), a protein variant where all WUS sequence except the homeodomain (HD), the (mutated) WUS box and the EAR motif have been replaced by flexible synthetic linker sequences, therefore representing a minimal WUS-like transcription factor. MiniMeΔbox has been reported to move to the L2 and L1 excessively and expression of transcriptionally active MiniMe protein results in massive over-proliferation of stem cells and drastic increase in meristem size ^2^. We also quantified distribution patterns of WUSCHEL RELATED HOMEOBOX 13 (WOX13) protein (WOX13-linker-FP), the most distant member of the gene family and the only *Arabidopsis* WOX gene not containing a WUS box ^32^. Together with the fact that WUS and WOX13 sequences outside the homeodomain are unrelated and the proteins are not functionally equivalent, WOX13 therefore represents a naturally evolved control for WUSΔbox.

When expressed from the *WUS* promoter, WUSΔbox was observed in the OC, as well as all cell layers above and showed the WUS-typical L3 to L1 gradient in the center of the SAM (Fig 2A). 2xGFP was similarly present in upper meristematic layers, with a shallow L3-L1 decreasing gradient, but since it was lacking an NLS, it did not appear distinctly nuclear. In contrast, 2xGFP-NLS showed exclusive nuclear localization, as expected from the addition of a strong NLS, and while consistently present in cells of the OC, it was only rarely observed in individual L2 cells and never in the L1. NES-WUSΔbox was mostly excluded from the nucleus, but otherwise exhibited a similar distribution pattern as WUSΔbox and 2xGFP. MiniMeΔbox on the other hand, excessively moved to apical meristematic layers, but also towards the periphery, and in fact the majority of protein seemed to be present in L2 and L1 even though the protein was expressed in the OC. WOX13, which showed nuclear localization similar to WUSΔbox, was distinctly present in the L2, but only a marginal amount of WOX13 protein could be observed in the L1.

**Figure 2.**
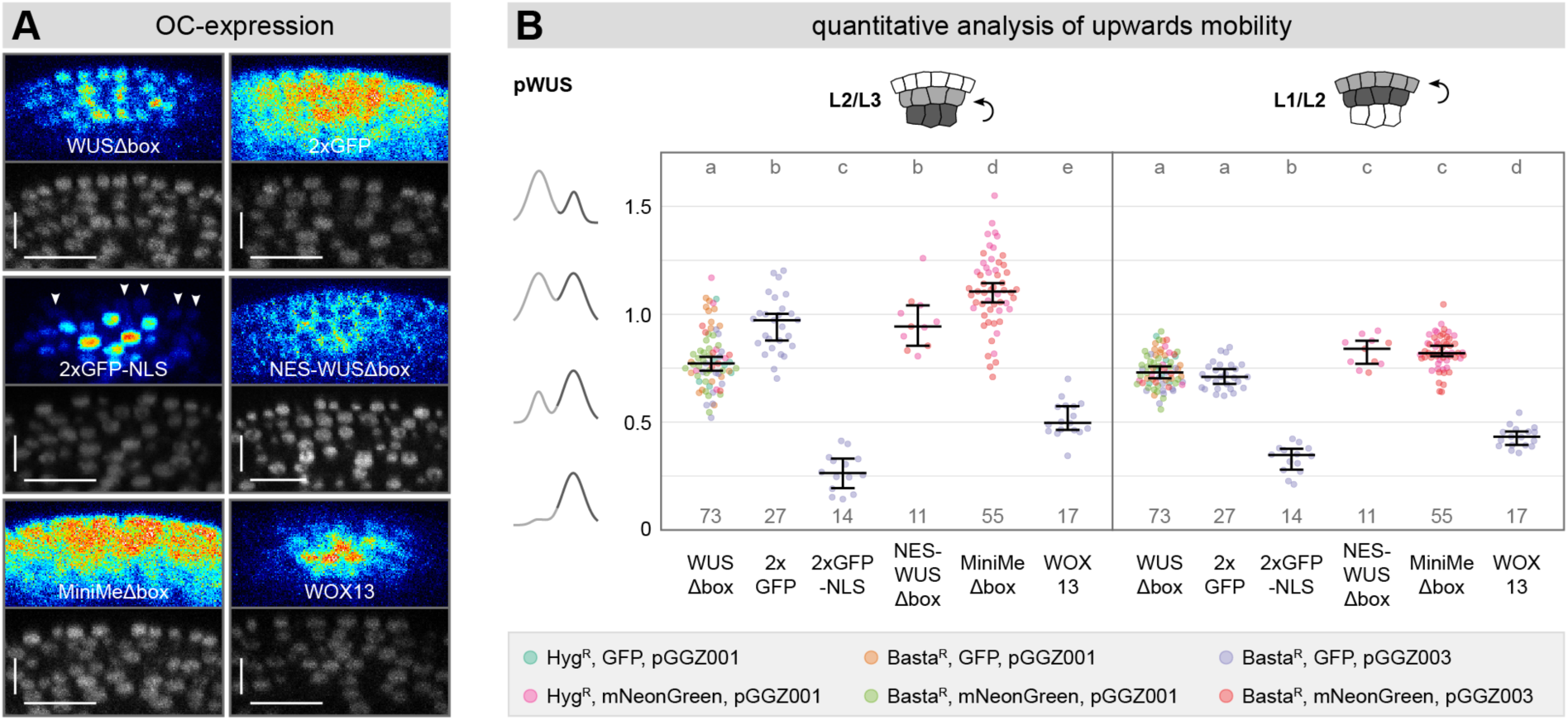
Analysis of basal to apical protein mobility in the shoot apical meristem. **(A)** Live-cell imaging of different fusion proteins expressed from the *WUS* promoter (T1 plants). All plants contained a genetically encoded, ubiquitous nuclear reporter (T3 line). Imaging was performed from a native top-view perspective. White arrows point out weak nuclear signal. Scale bars represent a length of 15 µm. **(B)** Quantitative analysis of upwards mobility for different fusion proteins expressed from the *WUS* promoter (T1 plants) using the ITQT tool. Different colors represent different construct designs. Lines represent population median values with a 95% confidence interval. Sample numbers are indicated below each population. Letters represent the result of pairwise comparisons by the ANOVA-TukeyHSD test (groups sharing a common letter are not significantly different at the 0.01 level).

Quantitative analysis of the images using ITQT (Fig 2B) supported our visual inspection: 2xGFP-NLS (L2/L3 median ratio 0.264, n = 14) barely showed any L2 signal, but moved slightly better than 3xGFP-NLS (data not shown), suggesting the protein was not completely immobilized but rather represents a diffusion control with a strong NLS. WOX13 (L2/L3 median ratio 0.497, n = 17) moved more efficiently from L3 to L2 than proteins tagged with a strong NLS, but significantly less than WUSΔbox (L2/L3 median ratio 0.772, n = 73), despite exhibiting a very similar cytoplasmic to nuclear fluorescence ratio. Interestingly, the addition of a nuclear export signal to a WUSΔbox fusion protein (NES-WUSΔbox) led to increased movement (L2/L3 median ratio 0.943, n = 11), not significantly different from 2xGFP (L2/L3 median ratio 0.972, n = 27). MiniMeΔbox (L2/L3 median ratio 1.105, n = 55) showed very high L3-to-L2 mobility, with the majority of plants within the population having a L2/L3 ratio >1, indicating that the amount of protein detected in the L2 was higher than in the L3 where the protein was synthesized. The difference between MiniMeΔbox and 2xGFP was statistically significant and only a small number of individual 2xGFP plants displayed a L2/L3 ratio >1. Considering that MiniMeΔbox tagged with a single fluorophore is slightly smaller than 2xGFP and both appeared similarly nuclear, it seemed plausible, at first glance, to assume that both proteins moved via passive diffusion and that the difference in mobility would result from the difference in size. However, passive diffusion could not explain the formation of a concentration maximum outside the domain of protein synthesis. We therefore propose that unlike for 2xGFP, MiniMeΔbox protein mobility is not only dependent on passive diffusion, but likely involves an unknown mechanism of active transport.

Looking at the movement from the L2 to the L1, we found more evidence for this hypothesis: Again, MiniMeΔbox (L1/L2 median ratio 0.820, n = 55) displayed a high layer ratio, moving significantly better than the 2xGFP diffusion control. In addition, WUSΔbox (L1/L2 median ratio 0.730, n = 73) and 2xGFP (L1/L2 median ratio 0.710, n = 27) showed no statistically significant difference, regardless of the fact that 2xGFP does not have a nuclear localization signal and hence faces less diffusion barriers than WUSΔbox. Importantly, both WUSΔbox and MiniMeΔbox efficiently bind chromatin, thus immobilizing a fraction of molecules, whereas for 2xGFP the entire pool is able to move from cell to cell. Consistently, NES-WUSΔbox (L1/L2 median ratio 0.840, n = 11), which was mostly excluded from the nucleus despite containing the WUS homeodomain, increasing the fraction of mobile, chromatin-unbound protein, showed significantly higher mobility than 2xGFP. In contrast, WOX13 (L1/L2 median ratio 0.432, n = 17), which unlike 2xGFP shares nuclear localization and chromatin binding with WUSΔbox and MiniMeΔbox, thus representing a second, likely more relevant control, moved significantly less from the L2 to the L1, compared to WUSΔbox, again supporting the notion of a protein-specific active transport component for WUS mobility.

To further investigate to what extend WUS mobility might be influenced by the cellular environment of sub-domains within the meristem, we expressed the same set of fusion proteins from the epidermal *ML1* promoter and analyzed their capacity for apical to basal movement from the L1 to the L3. Visual analysis (Fig 3A) revealed no striking differences compared to apical mobility from the OC. 2xGFP-NLS located almost exclusively in the cell layer where the fusion protein was expressed, while WOX13 reproducibly moved from the L1 to the L2, but hardly beyond. In contrast, WUSΔbox, NES-WUSΔbox, MiniMeΔbox and 2xGFP were observed not only in L1 and L2, but also in the L3, moving at least 3-4 cell layers downwards, qualitatively confirming previous findings ^2^. However, analysis of our imaging results by ITQT revealed significant differences (Fig 3B): Downwards mobility from the L1 to the L2 was low for 2xGFP-NLS (L2/L1 median ratio 0.236, n = 28) and WOX13 (L2/L1 median ratio 0.615, n = 23) moved more than NLS tagged fusion proteins, but significantly less than WUSΔbox (L2/L1 median ratio 0.829, n = 90). Again, WUSΔbox was not significantly different from 2xGFP (L2/L1 median ratio 0.805, n = 11), even though WUSΔbox was strongly nuclear localized. On the one hand this confirmed that WUSΔbox moves more than a diffusion control and on the other hand revealed that active WUS transport may facilitate protein movement also in the apical-basal direction, albeit less efficiently than upwards, as found in the *in-vivo* situation. NES-WUSΔbox (L2/L1 median ratio 1.004, n = 39) showed significantly higher mobility than WUSΔbox – in line with our previous data – but curiously also moved significantly more than MiniMeΔbox (L2/L1 median ratio 0.886, n = 40) for which a larger amount of fusion protein remained within the layer of synthesis compared to expression from the *WUS* promoter. This behavior was consistent with a reduced active mobility component towards basal layers and hinted to the fact that the responsible mechanism may be unable to promote mobility for nuclear proteins in the L1, whereas it able to do so in L3 and L2, potentially representing a way to enrich nuclear WUS within the stem cells.

**Figure 3.**
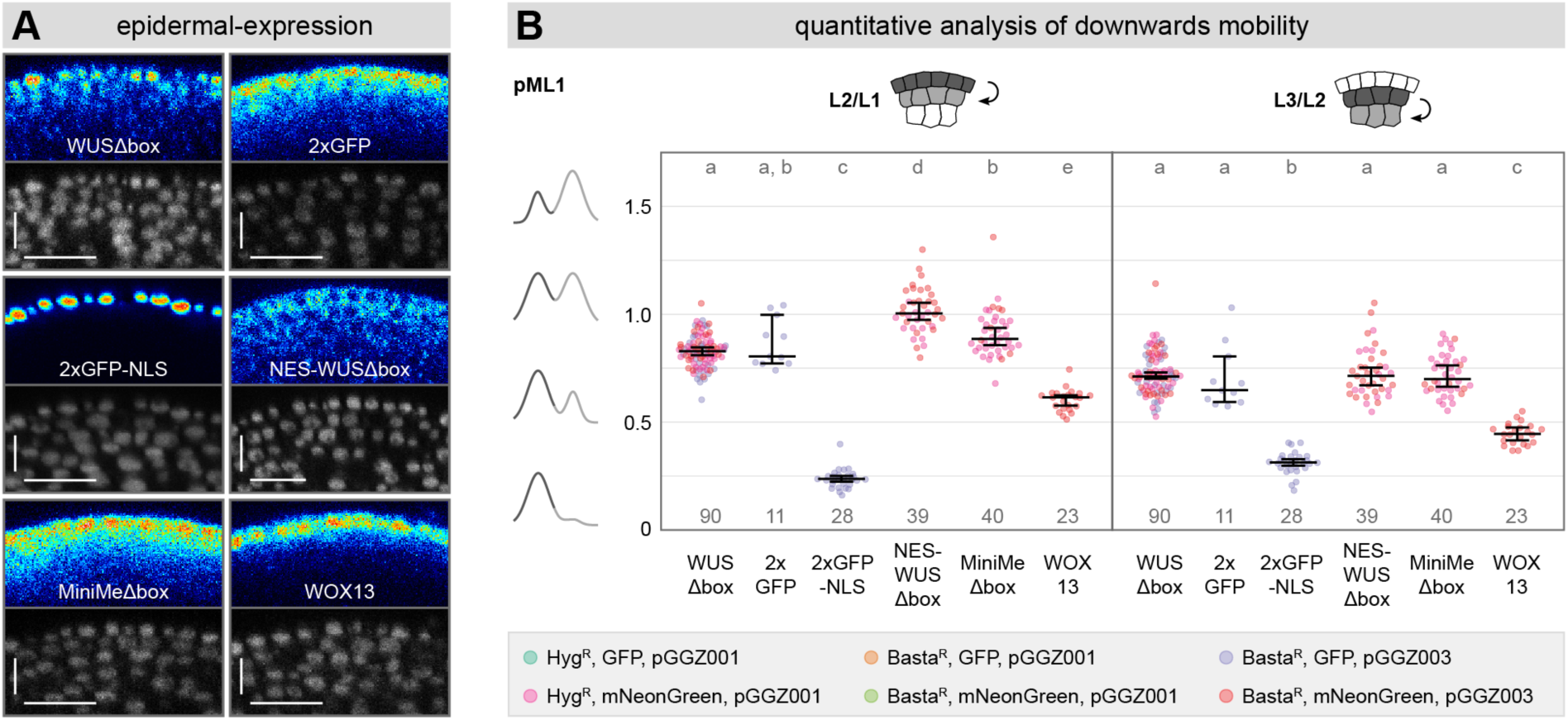
Analysis of apical to basal protein mobility in the shoot apical meristem. **(A)** Live-cell imaging of different fusion proteins expressed from the *ML1* promoter (T1 plants). All plants contained a genetically encoded, ubiquitous nuclear reporter (T3 line). Imaging was performed from a native top-view perspective. White arrows point out weak nuclear signal. Scale bars represent a length of 15 µm. **(B)** Quantitative analysis of downwards mobility for different fusion proteins expressed from the *ML1* promoter (T1 plants) using the ITQT tool. Different colors represent different construct designs. Lines represent population median values with a 95% confidence interval. Sample numbers are indicated below each population. Letters represent the result of pairwise comparisons by the ANOVA-TukeyHSD test (groups sharing a common letter are not significantly different at the 0.01 level).

When analyzing mobility from the L2 to the L3, MiniMeΔbox (L3/L2 median ratio 0.700, n = 40) moved better than 2xGFP (L3/L2 median ratio 0.648, n = 11), but slightly worse than WUSΔbox (L3/L2 median ratio 0.712, n = 90) and NES-WUSΔbox (L3/L2 median ratio 0.714, n = 39) and all four fusion proteins were not significantly different according to statistical analysis. For WOX13 (L3/L2 median ratio 0.445, n = 23), L2-to-L3 mobility was lower than for WUSΔbox and also lower than WOX13 L1-to-L2 mobility, in line with visual analysis of fluorescence images (Fig 3A).

Interestingly, when we considered the combined data for all layer transitions, especially regarding the comparison between WUSΔbox and 2xGFP, it became evident that even though WUS mobility is not directional per se, it appeared likely to be regulated in a layer specific manner: All ratios for mobility in adjacent cell layers were highly similar between WUSΔbox and 2xGFP, indicative of active transport of WUS to overcome the barriers imposed by chromatin binding and nuclear localization, except for the transition from L3 to L2. Here, movement of WUSΔbox was significantly lower compared to 2xGFP, suggesting the presence of a regulatory mechanism that restricts WUS mobility by retaining the protein in the OC. This conclusion was supported by the fact that MiniMeΔbox, which lacks important regulatory sequences ^2^, showed clear signs of active transport, such as forming a concentration maximum outside the domain of synthesis, but at the same time appeared devoid of retention signals, since its L3 to L2 ratio was not different from other cell-to-cell transitions. In contrast, NES-WUSΔbox behaved very similarly to WUS and therefore appeared to be subjected to the same regulatory mechanism, while WOX13, which is an evolutionary distant member of the WOX gene family, containing a homeodomain but sharing little general sequence homology with WUS, did neither appear to be actively transported nor affected in a layer-specific manner. Taken together, qualitative inspection of our data suggested that WUS mobility is facilitated by an unknown active transport mechanism, which is at least partially non-directional and is counteracted by protein retention in the L3.

In light of the quantitative nature, as well as the size and complexity of our dataset, including multiple proteins expressed from different promoters, we developed mechanistic mathematical models to systematically analyze different mobility scenarios and validate their plausibility using data-based model inference. The distribution of proteins in the SAM has been modeled previously in terms of differential equations describing dynamics of hormonal signaling ^33^, coupling them to organ growth ^34^ or at cellular resolution ^35^. Since ITQT analysis combines fluorescence signals of cells that belong to the same cell layer to a single value, we propose using the mathematical approach of ordinary differential equations which matches the resolution of our data and allows describing dynamics of different protein quantities following transport and degradation processes at different localizations (Fig 4A). As a result, our model depicts the meristem as three, linearly connected cellular domains, each representing one of the tissue layers L1, L2, L3 (where L1 connects to L2 and L2 connects to L3). Consequently, transitions from one layer to another can only occur between layers that are connected (from L1 to L2 or in the reverse direction; from L2 to L3 or in the reverse direction) and potential differences in signal distribution on a single-cell level along the central-peripheral axis are neglected, since they are also not captured by ITQT analysis. Additionally, the model distinguishes a cytoplasmic and a nuclear compartment for each cellular domain (= cell layer), to which proteins were proportionally allocated, based on known properties of the proteins and on visual observations from the imaging data. This allowed us to account for protein-specific differences in the cytoplasmic to nuclear ratio and for nuclear shuttling. Cell-to-cell translocation and protein degradation are only assumed for the pool of proteins currently present in the cytoplasmic compartment. We also tested our hypothesis for L3-specific regulation of WUS mobility by modeling protein retention specifically in the nuclear compartment of the L3. For the comparison of model simulation to the experimental data nuclear and cytoplasmic values were in the end combined into a single value per layer, to match the original ITQT output. Protein synthesis can take place either in the L3 or in the L1 to simulate expression from the *WUS* promoter or from the *ML1* promoter, respectively.

**Figure 4.**
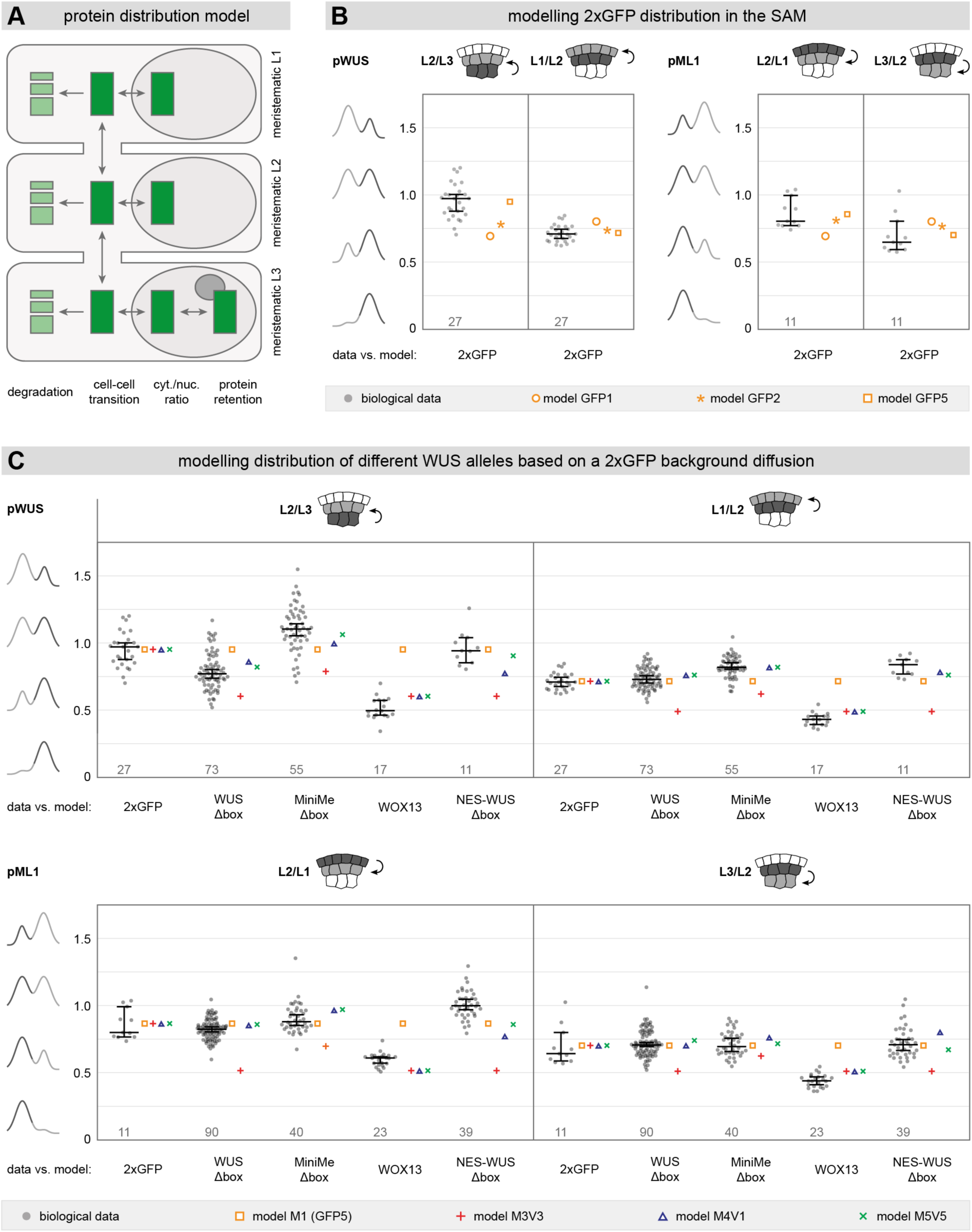
A mechanistic mathematical model for protein mobility the shoot apical meristem. **(A**) Schematic representation of a mechanistic modelling approach for protein distribution in L1, L2 and L3 of the SAM. The model takes into account degradation, cell-cell transition, cytoplasmic/nuclear protein ratio and protein retention in the L3. **(B**) Modeling of different scenarios for protein distribution of 2xGFP in the SAM. Data predicted by the model is compared to biological data acquired with ITQT analysis. Sample numbers of the biological data are indicated below each population. **(C**) Modeling of different scenarios for protein distribution of WUS fusion proteins in the SAM. Data predicted by the model is compared to biological data acquired with ITQT analysis. Sample numbers of the biological data are indicated below each population.

Mathematical models were built in an iterative manner; starting with the simplest set of assumptions and step-wise increasing the complexity of the model, each time validating the new model against experimental data. We started with a background mobility model, describing the behavior of 2xGFP to infer movement of a protein with similar size as WUS-linker-FP. Then, we modeled different scenarios testing various biologically motivated parameters for each layer, such as protein transport, degradation, or retention. First, we made basic parameter assumptions setting (1) constant protein degradation (2xGFP) in all layers; (2) constant and homogeneous (non-directional) diffusion/transport in/between the layers; (3) assuming a purely cytoplasmic protein; (4) without protein retention in L3. To our surprise, using these specifications we were unable to identify parameters that allowed the model to replicate the observed data of the *pWUS::2xGFP* and *pML1::2xGFP* datasets (Fig 4B). Regardless of the promoter used, we observed that 2xGFP showed a higher L2/L3 ratio together with a lower L1/L2 ratio than predicted by a diffusion model (model GFP1), indicating that *in vivo* 2xGFP accumulated in the L2. We then modified the model, allowing layer-specific, non-uniform degradation of 2xGFP (model GFP2). This assumption improved the agreement between biological data and modelled data to an extent, however, there was no reason to assume divergent layer-specific turnover of 2xGFP, which is not an endogenous protein in *Arabidopsis thaliana*. So instead, we tested the effects of allowing variable transport of 2xGFP across cell layers, rather than simple diffusion which constitutes for transport of a given rate that is the same in all directions (L3 to L2 = L2 to L3 = L2 to L1 = L1 to L2), while keeping protein degradation uniform (model GFP5). Under these assumptions, the model was able to closely replicate the biological data for both promoters. Due to its nature as a non-native protein with no endogenous function in *Arabidopsis thaliana* it seems unlikely that 2xGFP would be transported by a mechanism other than passive diffusion, so these results may instead point to an asymmetric barrier function between cell layers in the meristem, explaining why 2xGFP translocated more easily into cells of the L2 than out of L2. While the cellular mechanisms of this complex behavior remained unclear, this finding allowed us to define a realistic background model for undirected and passive cell-to-cell movement, which we applied to all further analyses (Fig 4C). From here on we assumed the mobility estimated for 2xGFP as passive diffusion across asymmetric barriers for L2 and used it as background model for all other protein species. In the next step, we tested whether the mobility of WUSΔbox, MiniMeΔbox and WOX13 could be explained by the passive diffusion across asymmetric L2 barriers exhibited by 2xGFP. Therefore, we simulated the distribution of the respective species across L1, L2 and L3 assuming that their mobility was identical to that of 2xGFP as quantified above. In addition, we assumed uniform protein degradation between protein species and within all cell layers (model M1). As suggested by the qualitative analysis, this was not the case and the model was unable to describe the behavior of WUS: Based on the background model, identical layer ratios for 2xGFP, WUSΔbox, MiniMeΔbox and WOX13 were predicted, failing to explain for the observed differences in the distribution patterns of these proteins. When fitting the model simultaneously to the data of 2xGFP, WUSΔbox, MiniMeΔbox and WOX13 the agreement between model prediction and data for 2xGFP decreased even further. Since WUSΔbox, MiniMeΔbox and WOX13 are nuclear proteins, we next included nuclear/cytoplasmic compartmentalization in our model, defining nuclear ratios based on visual observations from the imaging data (nucleus to cytoplasm: 2xGFP 3:7, WUSΔbox 9:1, MiniMeΔbox 7:3, WOX13 9:1) and again tested whether a diffusion model with asymmetric barriers could reproduce the protein distribution observed in our data. However, while different nuclear partitioning of 2xGFP, WUSΔbox, MiniMeΔbox and WOX13 led to differences in protein concentration between the different protein species, these effects were uniform across all cell layers. Consequently, they did not have an impact on the layer-to-layer ratios and therefore could not explain the protein distribution in our data. Next, we added differential protein stability to our model, which was represented by a degradation term. For a first test, we allowed the model to find the optimal degradation parameter for each protein, which was then applied to all layers. Adding degradation improved the model output substantially, but the parameters identified by the model did not reflect the biology of the molecules, since it predicted 2xGFP and WUSΔbox degrade at similar rates, MiniMeΔbox and NES-WUSΔbox at about half the rate, and WOX13 five times faster. While we do not have measured turnover rates for these proteins, we know from diverse experiments that 2xGFP is the most stable of the group, followed by and MiniMeΔbox, whereas WUSΔbox, NES-WUSΔbox and WOX13 are relatively short lived ^2^. We therefore set the degradation rates to reflect the biologically informed values (namely 2xGFP being four times as stable as WUSΔbox and MiniMeΔbox being twice as stable as WUSΔbox; WOX13 and NES-WUS have the same stability as WUSΔbox) and found that the fit of the model to the data was poor, suggesting that diffusion, nuclear localization and degradation within biologically meaningful ranges were not sufficient to explain the observed distribution patterns (model M3V3). We therefore included an additional process in the model, namely active transport, which was assumed to occur in addition to passive asymmetric diffusion. The consideration of this term was motivated not only by the poor performance of the model without it, but also by the experimental observation that MiniMeΔbox accumulated to higher levels outside the cells in which it was produced. Since we had shown that the mobility conferring regions of WUS are found inside the homeodomain ^2^, we assumed the active mobility component to be identical for WUSΔbox, NES-WUSΔbox and MiniMeΔbox, whereas for 2xGFP and WOX13 this parameter was fixed to 0. Including the active mobility component in the model significantly improved the overall fit to the experimental data for all proteins studied (model M4V1). Interestingly, the model deviated most from the experimental data for the transitions between the L3 and L2 with WUSΔbox mobility overestimated and MiniMeΔbox underestimated. This was also reflected in the derived parameters that showed a lower transport component for transitions between the L3 to L2 than between L2 and L1, whereas the values were fairly similar regardless whether apical or basal mobility was analyzed. Consistent with the previous result indicating heterogeneous translocation rates, assuming identical parameters for a given direction or layer transition made the model perform worse. Finally, to address the mechanisms that lead to reduced mobility of WUS and its derivatives from the L3, we added an L3 retention parameter to the model. In biological terms this would represent, for example, a protein expressed in the OC whose interaction with WUS would make the complex too large for translocation to the L2. As we had previously shown that sequences restricting WUS mobility are located in the unstructured parts outside the conserved domains, we restricted the retention to WUSΔbox and NES-WUSΔbox, the only proteins containing the relevant parts. Indeed, the addition of an L3 specific retention term further improved the model fit to the experimental data, even when all transport terms were retained from the previous version. Importantly, the model predictions further improved significantly, after we allowed the transport terms for the transitions between L3 and L2 to be optimized in the presence of retention (model M5V5). The final model was able to robustly predict results for all proteins tested from the WUS promoter, but showed some deviations from the experimental data for NES-WUSΔbox and WOX13 expressed from the ML1 promoter. Importantly, the model revealed that proteins containing the WUS homeodomain require a much higher transport term compared to 2xGFP to explain their steady state distribution patterns. The model predicted the active component to represent roughly 70% to 80% of the total mobility depending on the tissue layer. The deviation of NES-WUSΔbox and WOX13 expressed from the ML1 promoter could either be due to technical limitations in our data, or to the fact that the L1 represent the border layer to the environment from which only basally oriented mobility is possible. Hence, we can conclude that active transport of the WUS homeodomain occurs similarly across all layers and directions.

Taken together, systematic mathematical modeling using biologically meaningful assumptions on the model parameters and a data-based quantitative model validation suggested that steady state WUS protein distribution in the SAM is the result of a complex interplay of asymmetric diffusion, active transport and protein retention. Importantly, it became clear that active transport appears to be required in all cell layers and for both the apical, as well as basal direction. In contrast, protein retention appears to be specific to the OC, arguing for WUS to be trapped in a protein complex with partners specific to the L3, such as HAM1 or HAM2 ^24–26^. In contrast, WOX13-GFP consistently moved less than 2x-GFP in line with its nuclear localization and ability to bind to chromatin. Since quantitatively, the difference between 2xGFP and WOX13-GFP was very similar for all cell-to-cell transitions, the results of the model were compatible with the hypothesis that WOX13-GFP distribution is the result of diffusion and therefore represents the behavior of a control transcription factor.

After having delineated the mechanisms for active WUS transport in the center, to facilitate movement from the OC to the CZ along the apical-basal axis, we next asked whether there were any differences in protein mobility between the central zone and the periphery. Therefore, we analyzed the behavior of selected fusion proteins expressed from the *ML1* promoter – whose activity is even throughout the meristematic epidermis – and extended our analysis to a hollow cylinder (dognut) with a diameter of 40 µm and a ring thickness of 10 µm, placed around the 20 µm cylindrical column in the center of the meristem (Fig 5A). However, our quantitative analysis did not reveal any statistically significant difference in downwards mobility between the center and the periphery for WUSΔbox, NES-WUSΔbox and MiniMeΔbox (Fig 5B), suggesting that the mechanism for WUS transport is equally active in the CZ as well as in the PZ. Similarly, WOX13 and 2xGFP did not show differential mobility between the center and the periphery (Fig 5B), suggesting that (downwards-directed) cellular connectivity is also unlikely to be differentially regulated between CZ and PZ.

**Figure 5.**
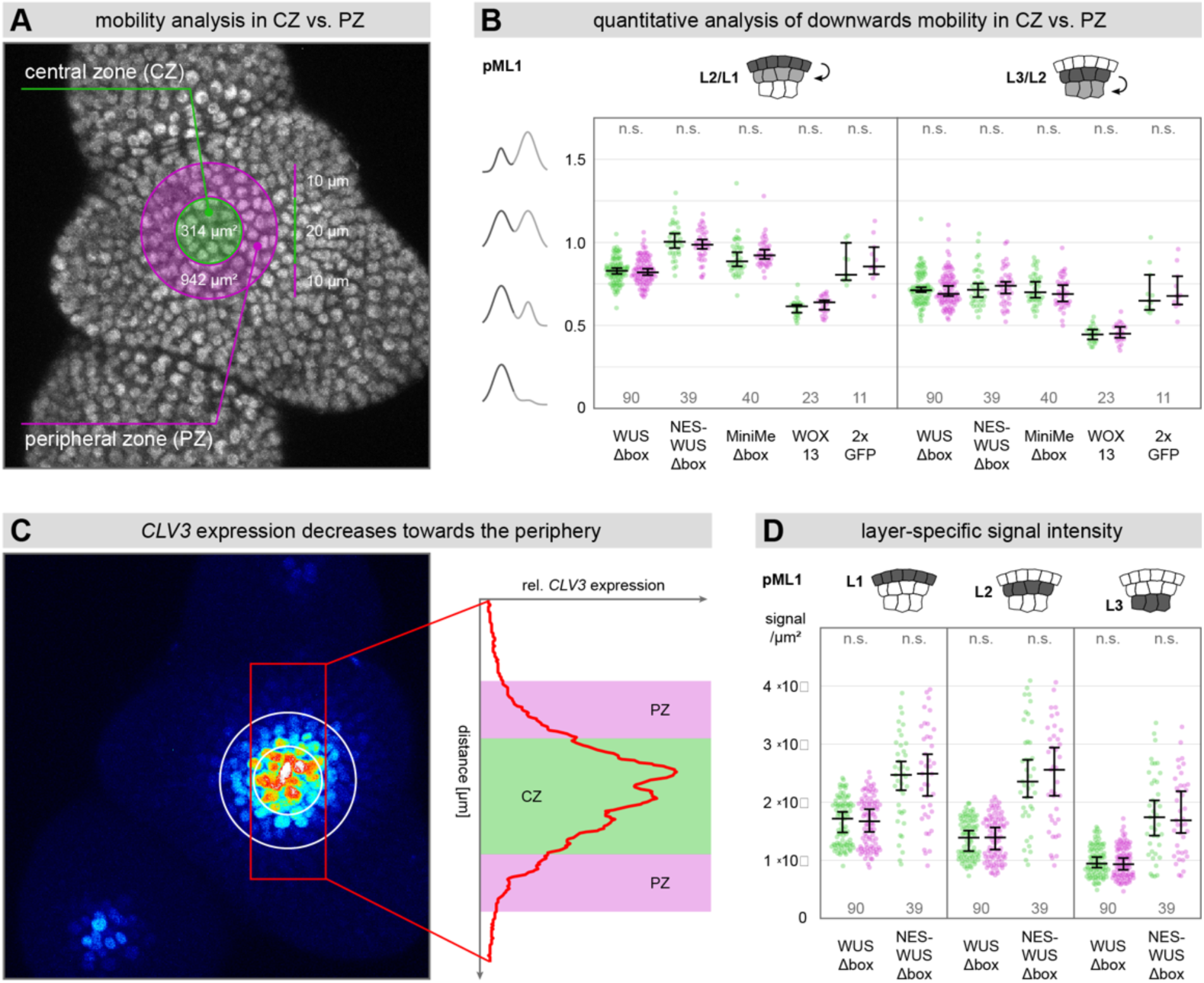
Analysis of differential WUS mobility between CZ and PZ. **(A)** Representation of ITQT-analysis domains covering CZ (green) and PZ (lilac) in the shoot meristem. **(B)** Quantitative analysis of downwards mobility for different fusion proteins expressed from the *ML1* (T1 plants) using the ITQT tool in CZ vs. PZ. Green data points represent CZ data; lilac data points represent PZ data. Lines represent population median values with a 95% confidence interval. Sample numbers are indicated below each population. Data was analyzed with a Student’s T-test (n.s. = no significant difference = p < 0.05). **(C)** Representation of relative CLV3 expression in CZ (green) vs. PZ (lilac). **(D)** Quantification of layer-specific signal intensity for nuclear and cytosolic WUS (WUSΔbox and NES-WUSΔbox) in CZ (green) vs. PZ (lilac). Signal intensity was normalized to intensity/µm^2. Lines represent population median values with a 95% confidence interval. Sample numbers are indicated below each population. Data was analyzed with a Student’s T-test (n.s. = no significant difference = p < 0.05).

A recent study based on over-expression studies of a WUS fusion with the glucocorticoid receptor (GR) and GFP along with peptide treatments has hypothesized that the CLV3 peptide controls nuclear retention of WUS and its stability in the cytoplasm ^23^. Our dataset and analysis tool now allowed us to test different elements of this hypothesis by comparing the behavior of two WUS-GFP alleles that differ in their nuclear localization in parts of the meristem that have either high or low levels of endogenous *CLV3* expression, respectively (Fig 5C). Therefore, we analyzed layer-specific fluorescence intensity, as a measure for absolute protein levels, for both WUSΔbox (nuclear) and NES-WUSΔbox (cytoplasmic) in the center (high *CLV3*) and periphery (low *CLV3*) of the SAM. If stability of the WUS protein was dependent on CLV3, whose expression levels decrease from the central zone towards the periphery and from the L1 to the L3, we would expect to see differential accumulation of WUS protein between the center and the peripheral zone. However, protein levels for WUSΔbox, which is mostly nuclear localized, as well as for NES-WUSΔbox, which localizes predominantly in the cytoplasm, did not differ between CZ and PZ in either L1, L2 or L3 (Fig 5D). Thus, our experiments leveraging the natural CLV3 gradient and using WUS fusions much more similar to endogenous WUS than GFP-WUS-GR, did not support a role for the CLV3 peptide in controlling WUS stability in the cytoplasm or nucleus.

Having provided strong evidence for the presence of an active transport mechanism that facilitates cell-to-cell movement of WUSΔbox, NES-WUSΔbox and MiniMeΔbox, but not of WOX13, we were curious to find out how specificity of active transport could be achieved. Because the super moving MiniMeΔbox only contains the WUS homeodomain, the (mutated) WUS box and the EAR motif, whereas the low moving WOX13 only shares the mildly conserved homeodomain, but neither has WUS box nor EAR motif, we decided to focus on these domains. To test whether the EAR motif might mediate active WUS transport, we created a fusion protein where both the WUS box as well as the EAR motif were mutated (WUSΔboxΔEAR) ^31^. We again used the *WUS* promoter to drive expression and performed life-cell imaging from a top-view perspective on T1 populations. Visual inspection of WUSΔboxΔEAR(-linker-FP) showed no apparent differences to WUSΔbox: Both proteins displayed the same L3-to-L1 decreasing gradient and showed similar, strong nuclear localization. Furthermore, quantitative analysis of vertical protein distribution using ITQT revealed no statistically significant difference between WUSΔboxΔEAR (L3/L2 median ratio 0.798, L2/L1 median ratio 0.704, n = 24) and WUSΔbox (L3/L2 median ratio 0.772, L2/L1 median ratio 0.730, n = 73) (Fig 6A), suggesting that the EAR motif is unlikely to control mobility or stability of the WUS protein. Next, we wanted to revisit the role of the WUS-box leveraging the sensitivity afforded by ITQT. We therefore compared the distribution of transcriptionally active WUS-linker-GFP in *wus* mutants to inactive WUSΔbox and WUSΔboxΔEAR in wild-type meristems. Visual inspection suggested that the inactive proteins were slightly less nuclear than active WUS, likely reflecting their inability to bind nuclear interaction partners, such as the transcriptional repressor TOPLESS (TPL) ^36,37^, however, there was no qualitative difference in protein distribution between the three forms of WUS, confirming previous findings ^2^. Quantitative analysis using ITQT showed that wild-type WUS moved significantly less from the L3 to the L2 as well as from the L2 to the L1 compared to versions of WUS with mutations in the WUS-box. These findings were in line with the important function of the WUS-box in mediating the interaction with the transcriptional repression machinery and likely reflect the fact that WUS molecules embedded into this large protein complex are unable to move from cell to cell. Interestingly, for L3-to-L2 mobility, the difference between functional and mutated versions of WUS was small, supporting our hypothesis of important regulation via protein retention in the L3, that overrides the effect of trapping in a large protein complex and which is independent of the WUS-box or EAR motif. Therefore, while the WUS-box contributes to WUS retention, we propose that between the WUS homeodomain, the WUS box and the EAR motif, the homeodomain is a driver for protein mobility, with yet other sequences playing a mayor regulatory role, in line with previous findings ^2^.

**Figure 6.**
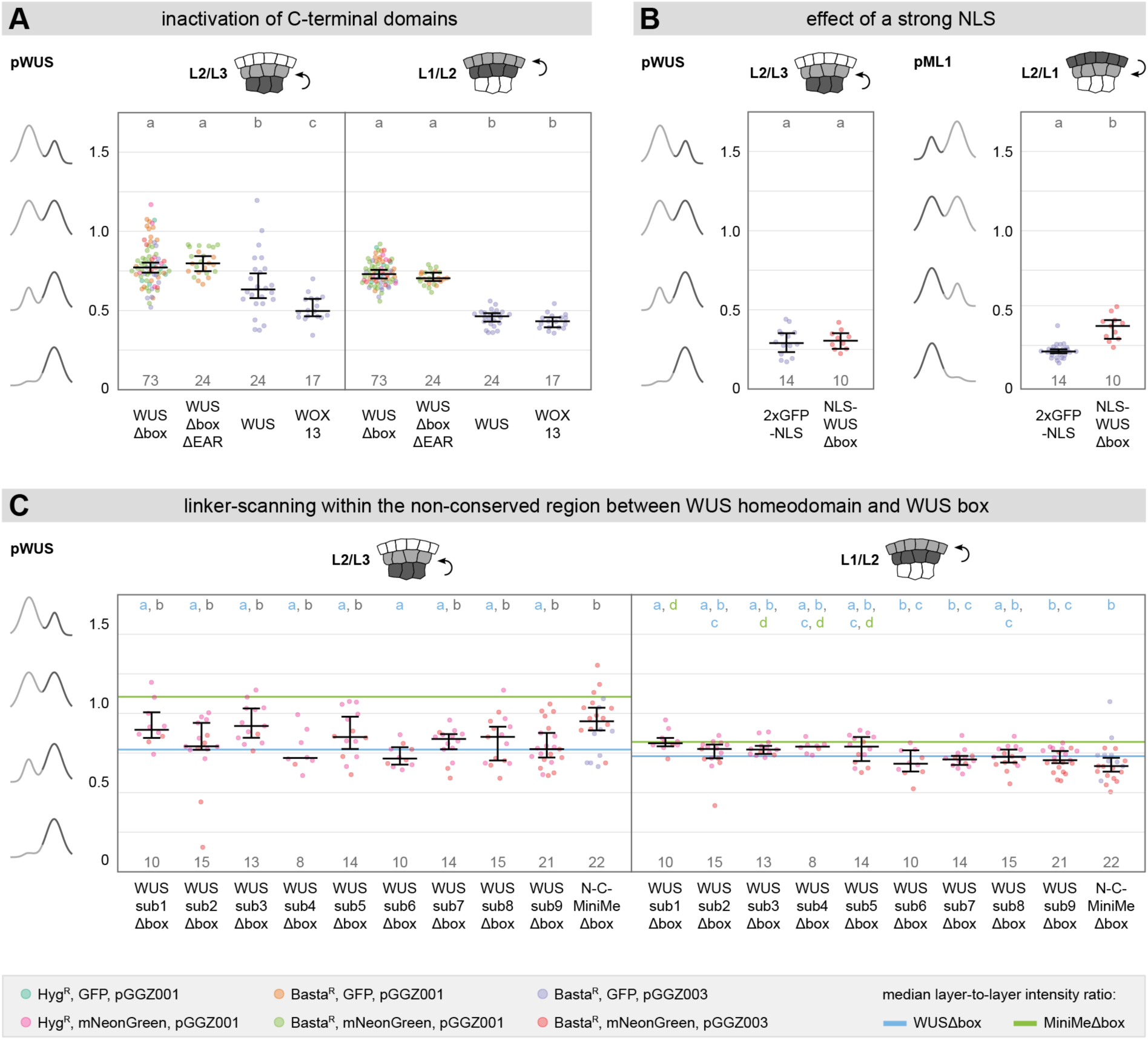
Role of WUS C-terminal amino acid motifs for mobility. **(A)** Quantitative analysis of the effects of WUS C-terminal domains on upwards mobility using the ITQT tool. Different colors represent different construct designs. Lines represent population median values with a 95% confidence interval. Sample numbers are indicated below each population. Letters represent the result of an ANOVA-TukeyHSD statistical test (significance value = 0.01; same letters indicate no statistically significant difference). **(B)** Quantitative analysis of the effects of a strong NLS on upwards mobility using the ITQT tool. Different colors represent different construct designs. Lines represent population median values with a 95% confidence interval. Sample numbers are indicated below each population. Letters represent the result of an ANOVA-TukeyHSD statistical test (significance value = 0.01; same letters indicate no statistically significant difference). **(C)** Quantitative analysis of upwards mobility for different linker-scanning fusion proteins expressed from the *WUS* promoter (T1 plants) using the ITQT tool. Different colors represent different construct designs. Lines represent population median values with a 95% confidence interval. Sample numbers are indicated below each population. Letters represent the result of pairwise comparisons by the ANOVA-TukeyHSD test (groups sharing a common letter are not significantly different at the 0.01 level).

Next, we asked whether the mechanism for active WUS transport was strong enough to overcome the limitations effectuated by the addition of a strong nuclear localization signal. To this end, we compared 2xGFP-NLS with NLS-WUSΔbox(-linker-FP), specifically with regard to the transition from the cell layer of expression to the first non-expressing layer (L2/L3 ratio for *pWUS*; L2/L1 ratio for *pML1*). When we looked at protein mobility in upwards direction from the L3 to the L2, we saw no significant difference between 2xGFP-NLS (L2/L3 median ratio 0.264, n = 14) and NLS-WUSΔbox (L2/L3 median ratio 0.279, n = 10), which seemed to suggest that active WUS transport was unable to overcome the effect of a strong NLS. Interestingly, however, downwards, from the L1 to the L2, NLS-WUSΔbox (L2/L1 median ratio 0.398, n = 11) moved significantly better than 2xGFP-NLS (L2/L1 median ratio 0.236, n = 28), not only showing that NLS-WUSΔbox is actively transported despite containing a strong NLS, but providing further evidence for the regulation of active transport via protein retention in the L3.

To further investigate the question of how active WUS transport is regulated, we again used our knowledge of the WUSΔbox and MiniMeΔbox fusion proteins and performed a systematic linker-scanning experiment (Fig S3). Our first experimental allele was N-C-MiniMeΔbox, a hybrid protein that contained the endogenous WUS N-terminus and C-terminus but has all sequence between the homeodomain and the (mutated) WUS box replaced by the same linker stretch used in the MiniMe protein. In addition, we created nine substitution alleles by replacing adjacent 17 amino acids (aa) sequence stretches of WUSΔbox located between homeodomain and WUS box with unspecific linker-sequence. All WUS variants were C-terminally tagged, using linker-GFP or linker-mNeonGreen, and expressed from the *WUS* promoter for comparison to WUSΔbox and MiniMeΔbox Interestingly, when we analyzed movement from the L3 to the L2 (Fig 6C), N-C-MiniMeΔbox (L2/L3 median ratio 0.950, n = 22) moved more than WUSΔbox but less than MiniMeΔbox and was significantly different from both, suggesting the presence of additional regulatory sequences within the N-or C-terminus. Analysis of the linker-substitution alleles revealed that while several substitutions in the central part of WUS seemed to have increased protein mobility, no single version was able to reproduce the dramatic increase observed for MiniMe(Δbox) in this study or in earlier work ^2^. All substitutions, except for WUSsub6Δbox, which was significantly different from N-C-MiniMeΔbox but not from WUSΔbox, were not different from both WUSΔbox as well as N-C-MiniMeΔbox, indicating slightly increased, intermediate mobility. For L2-to-L1 mobility, all fusions proteins showed similar ratios: Statistical analysis grouped all substitution alleles as well as N-C-MiniMeΔbox together with WUSΔbox and some substitutions were additionally grouped with MiniMeΔbox, further supporting the hypothesis that regulation of active WUS transport is mostly a property of the organizing center in the L3.

Taken together, in this study we present evidence that WUS transport from the organizing center to the stem cells in the central zone is facilitated by an active transport mechanism. Our data further suggests that the WUS homeodomain is necessary for active transport and might mediate specificity of the process. We propose that this mechanism is non-directional and not exclusive to sub-domains within the meristem, but is controlled via protein retention in the L3, which in turn is dependent on WUS protein sequence within the N-or C-terminus on the one hand and the sequence stretches between the homeodomain and the WUS box on the other hand.

## Discussion

Starting with its initial description based on a meristematic phenotype in the model plant *Arabidopsis thaliana* roughly 25 years ago ^28^, *WUS* has attracted the interest of generations of plant researchers and continues to do so until today. As a homeodomain transcription factor ^1^, WUS sits at the center of multiple regulatory feedback loops ^8,9,14,19,38,39^, takes on the capacity of a transcriptional activator or repressor depending on tissue context ^1,4,31,32,36,39,40^, and is indispensable for stem cell maintenance in shoot and floral meristems ^1^, floral patterning ^39^ and ovule development ^41^. In the shoot apical meristem, WUS protein acts as a mobile factor to connect niche and stem cells by moving from the organizing center in the L3 to the cells in the epidermis via plasmodesmata ^2,3^. How this cell-to-cell mobility is achieved, how WUS levels in the SAM are regulated and how the robustness of the WUS gradient is maintained has been a matter of debate and a number of studies have reached partially contradictory conclusions, mostly based on the analysis of fluorescent images with tagged WUS protein, either via visual inspection ^2,3,21^ or simple image quantifications ^22,23^. We have previously discussed conflicting data and have pointed out the need for a systematic comparison of different WUS-tagging strategies using high-quality life-cell imaging and reliable, large-scale quantitative analyses ^42^. In this study, we tested four different transgenes, containing N-and C-terminal fusions between WUS and GFP, with and without a flexible serine-glycine linker, respectively, driven by identical *WUS* promoter (4.4 kb) and terminator (2.8 kb) fragments for expression control and assembled with minimal cloning scars. We used these transgenes to generate homozygous, single-insertion lines complementing a *wus* null mutation and compared the fusion proteins with regard to rescue efficiency and *in-vivo* protein behavior. We found that N-terminally tagged WUS without linker (GFP-WUS) and C-terminally tagged WUS with linker (WUS-linker-GFP) were nearly indistinguishable: Both fusion proteins showed strong nuclear localization as well as the WUS-typical L3-to-L1 protein gradient ^2,3^ and efficiently rescued the *wus* mutation with low occurrence of meristematic phenotypes. C-terminal tagging without a flexible linker (WUS-GFP) on the other hand slightly increased protein mobility, had no effect on subcellular localization, but had a lower rescue efficiency with a high rate of meristem termination in the vegetative state. The addition of a flexible linker at the N-terminus (GFP-linker-WUS) increased protein mobility even more, while at the same time the protein showed a decrease in nuclear localization as well as the lowest rescue efficiency with a high rate of meristem termination in the reproductive state. We suspect that the C-terminal addition of the fluorophore interfered with the function of important C-terminal protein domains, while the N-terminal linker in this case provided too much freedom of movement for the fluorophore, allowing it to interfere with some functions of the N-terminal homeodomain. It is however interesting that both these fusion proteins showed phenotypes occurring at different developmental stages, suggesting that there may be specific roles of the homeodomain and WUS C-terminal domains in reproductive and vegetative development, respectively.

Since purely visual analysis of subtle quantitative differences, spanning several cells within a complex three-dimensional tissue such as the shoot apical meristem, can only detect qualitative differences, we developed the “Image Transformation and Quantification Tool’ (ITQT) for semi-automated alignment of meristematic cells and fluorescence signal quantification along the apical-basal axis. Using ITQT, we quantified the vertical distribution in upper meristematic cell layers for all four WUS fusion proteins. In line with our data on rescue efficiency, we found that GFP-WUS and WUS-linker-GFP were not significantly different from each other. Importantly, the comparatively lower accumulation of WUS-linker-GFP protein in L1 and L2, clearly contradicts the suggestion by Rodriguez and colleagues that the WUS-linker-GFP fusion represents an artificially stabilized protein variant ^21^.

In contrast, for WUS-GFP and GFP-linker-WUS, which both showed meristematic phenotypes at high frequency, suggesting impaired protein function or disturbed feedback regulation, vertical protein distribution was shifted upwards. Interestingly, an increase of WUS protein within *CLV3* positive cells should lead to over-proliferation ^8^, rather than the stem cell termination phenotypes observed, suggesting that these alleles may have reduced functionality. Therefore, we hypothesize that the observed increase in the amount of fusion protein in L1 and L2 may represent a mechanism to compensate for the partial loss of WUS function.

In general, we noticed that the variability of layer-to-layer intensity ratios was high, even when analyzing stable, non-segregating lines and further increased with distance from the epidermis. This observation seems plausible, since first, the circle-points for polar-transformation based ‘unrolling’ of meristem curvature were placed at the surface of the tissue and precision of the transformation is likely to decrease with increasing distance from its reference points. Second, the structure of the meristem becomes less ordered in deeper layers, due to periclinal cell divisions and increasing cell sizes, resulting in less precise alignment of cells that belong to the same layer even after polar transformation. To partially compensate for high variability and to increase statistical power, we decided to quantify T1 populations rather than established lines and pool results from multiple constructs carrying the same fusion protein. While this strategy will increase variability overall, it allowed us to control for effects of individual transgenic lines.

Miss-expression of functional WUS ^7^ and modifications to the WUS protein that shift its distribution upwards ^2^ lead to ectopic initiation of stem cells and over-proliferation. Conversely, blocking WUS movement from the OC ^2^ or *CLV3*-specific degradation of WUS protein ^7^ result in stem cell loss and meristem termination. To avoid changes in meristem structure resulting from such alterations, we used WUS fusions, which due to a mutation of their WUS box (Δbox) have no transcriptional activity ^31^. We expressed transcriptionally inactivated Δbox protein fusions from the *WUS* promoter as well as from the epidermal *ML1* promoter and quantified protein distribution along the apical-basal axis within the center of the meristem. We developed a mathematical modeling approach which allowed us to systematically compare different hypotheses about the mobility of the considered protein species. The models account for diffusion, active transport, protein retention as well as for the distribution of the molecules between nucleus and cytoplasm. We found strong evidence for an active transport mechanism that is specific to WUS and facilitates protein mobility regardless of the directionality in the shoot meristem: MiniMeΔbox, an artificial, mimimal, WUS-like repressor protein, in 78% of the plants (43/55) showed the formation of a concentration maximum outside of its expression domain in the OC (*WUS* promoter). For 2xGFP, which we used as a diffusion control, this occurred in only 33% of the plants (9/27), suggesting that the effect cannot only be explained from potential analysis artifacts, but likely is biologically relevant. Therefore, we hypothesized that MiniMeΔbox, unlike 2xGFP, is subject to a, potentially WUS specific, active transport mechanism. We tested this hypothesis using a series of mathematical models of increasing complexity. The mathematical modeling suggests that 2xGFP is subjected to passive diffusion with an asymmetric L2 barrier. The distributions observed for WUSΔbox and MiniMeΔbox can be explained by active transport which acts in addition to the passive asymmetric diffusion. Differences of the L2/L3 ratios for WUSΔbox and MiniMeΔbox can be captured if we assume that, in addition to active transport and passive diffusion WUSΔbox is retained in L3 by binding to nuclear proteins. We found more evidence for this hypothesis, when we compared the distribution of 2xGFP and WUSΔbox: Even though WUSΔbox is located in the nucleus and is able to bind chromatin with high affinity, both proteins showed the same L1/L2 ratio, suggesting an active mechanism compensating for the apparent diffusion barriers of WUSΔbox. Consequently, WUSΔbox tagged with a nuclear export signal (NES-WUSΔbox), preventing nuclear accumulation and likely the majority of chromatin binding events, showed L2 to L1 mobility significantly higher than WUSΔbox and 2xGFP, but comparable to MiniMeΔbox. On the other hand, WOX13, which also binds chromatin but despite being a member of the WUS-homeobox containing (WOX) family shares little sequence homology with WUS, in general as well as in the homeodomain, and does not contain a WUS box, moved significantly less than WUSΔbox, NES-WUSΔbox, MiniMeΔbox, suggesting that the former move via a WUS-specific, active transport mechanism, but also less than 2xGFP, mirroring the differential diffusion barriers between the two proteins not actively transported. Analysis of the same set of fusion proteins expressed from the epidermal *ML1* promoter supported the presence of WUS-specific, active transport, and additionally suggested that such a mechanism might not be limited to facilitating unidirectional, upwards mobility, since WUSΔbox, MiniMeΔbox and NES-WUSΔbox showed higher L1-to-L2 mobility than 2xGFP. Interestingly, the efficiency of downwards transport seemed lower than the efficiency of upwards transport, as evident from the comparison of layer ratios as well as visual impression, and unlike for upwards mobility, downwards, NES-WUSΔbox moved more than MiniMeΔbox. These differences might reflect the role of WUS as a transcription factor that binds open chromatin in L1 stem cells. It is conceivable that in L1 nuclear WUS, but not cytoplasmic WUS is prevented from leaving the stem cells in the center again. Alternatively, it is possible that in L1 there are more open binding sites for WUS, which would affect the mobility of WUSΔbox and MiniMeΔbox greatly and the mobility of NES-WUSΔbox only marginally. Analysis of mobility from L2 to L3 showed no statistically significant differences of WUS-like proteins compared to 2xGFP, however, 2xGFP, despite small sample size, showed the same tendency for lower mobility and had the lowest median ratio. In conclusion, these data supported our hypothesis that WUS mobility is facilitated via a partially bi-directional, WUS-specific, active transport mechanism. Our findings that adding an NES only caused subtle differences to the mobility of WUS along with our insights from modelling clearly suggest that the cytoplasmic to nuclear ratio of WUS has very little influence on its tissue distribution, in contrast to claims by Rodriguez and colleagues ^21^. This is also in line with the fact that WUS distribution is only analyzed at the steady state level and, therefore, the cytoplasmic to nuclear ration could only have an influence if the transition between nucleus and cytoplasm was much slower that the transition of WUS between cells.

The comparison of our side-view imaging data on the distribution of functional WUS within the WUS-linker-GFP rescue line to the top-view imaging data of a WUSΔbox-linker-FP T1 population, showed that WUS containing a functional WUS box moved less than WUSΔbox. This additional reduction in WUS mobility, compared to WUSΔbox, likely reflects the inability of Δbox proteins to interact with transcriptional co-factors, such as TPL ^36,37^, providing more freedom of movement for Δbox proteins. Interestingly though, functional WUS still moves better (L3 to L2) or similar (L2 to L1) to WOX13, even though WOX13, just like WUSΔbox, does not contain a WUS box, providing further evidence for active transport that is specific to WUS and does not apply to either 2xGFP or WOX13.

Specificity of active WUS transport seems to be exerted by the WUS homeodomain, that differs from the WOX13 homeodomain. MiniMeΔbox, which has lost all sequence except the homeodomain, the mutated WUS box and the EAR motif showed mobility strongly suggestive of active transport and a WUSΔboxΔEAR fusion protein moved indistinguishable from WUSΔbox, which we have also shown to likely move via an active mechanism.

Interestingly, despite reaching a different conclusion, Rodriguez and colleagues have previously presented data that seems to supports our hypothesis of active movement via the WUS homeodomain: When they expressed the first 134 amino acids of WUS, including the homeodomain, but not the C-terminal transcriptional domains, as an N-terminally tagged GFP fusion, the authors report ‘an intense and relatively uniform distribution of fluorescence in all cell layers’ ^21^ and indeed the protein showed strong upwards mobility and was prominently present in L1, L2 and L3. Their data seems to indicate little difference in concentration between the three uppermost cell layers, however this needs to be interpreted with caution, since the image shown is over-saturated and not suitable for more quantitative analysis. In any case, these results are reminiscent of our own data on MiniMeΔbox mobility, as well as of qualitative data on MiniMe(Δbox) mobility presented earlier ^2^.

It has also been hypothesized that the WUS gradient is regulated by nuclear-cytoplasmic partitioning and directly dependent on CLV3 peptide levels ^21,23^. When we analyzed the distribution of nuclear WUS (WUSΔbox) and cytoplasmic WUS (NES-WUSΔbox) in the center (high endogenous CLV3) versus the periphery (low endogenous CLV3), we did not observe any difference in layer-specific mobility or in protein levels between CZ and PZ. Similarly, the comparison of mobility ratios for 2xGFP and WOX13 between the center and the periphery did not suggest any differences in intercellular connectivity. From these results we can deduce that the endogenous CLV3 peptide does not contribute to the regulation of WUS mobility in contrast to the hypothesis put forward by Plong and colleagues, which relied on exogenous application of high levels of synthetic CLV3.

Instead, our data points towards a different mode of regulation based on protein retention in the L3, likely via protein-protein interactions: WUSΔbox moved similar to 2xGFP, despite its nuclear localization and ability to bind chromatin, suggestive of active transport, for all layer transitions, except for the transition from the L3 to the L2, where it moved significantly less compared to 2xGFP. Similarly, the mobility difference between WUS and WUSΔbox was smaller for the L2-to-L1 transition compared to the L3-to-L2 transition, suggesting an additional L3-specific reduction in mobility independent from the interaction with co-repressors via the WUS box. Interestingly, neither tagging with a strong NLS nor the addition of an NES seemed to negatively affect this regulation. However, L3-specific regulation did not apply to MiniMeΔbox, missing all endogenous WUS sequence except the homeodomain, the (mutated) WUS box and the EAR motif, similar to WOX13. Since a previous study, using qualitative analysis of the MiniMe protein, has defined regulatory sequence in the sequence stretch between the WUS homeodomain and the WUS box ^2^, we performed a linker-scanning experiment specifically within this region. To this end, we created fusion proteins where short stretches of 17 amino acids were replaced by serine-glycine linker, expressed them from the *WUS* promoter and quantified the vertical distribution of the proteins. Interestingly, no single substitution allele was able to recapitulate the severe mobility phenotype of the MiniMe(Δbox) protein. Instead, a number of alleles showed intermediately increased mobility from L3 to L2, compared to WUSΔbox and MiniMeΔbox. The same was true for N-C-MiniMeΔbox, a MiniMeΔbox allele where the N-terminus and the C-terminus have been re-introduced, suggesting that regulation of WUS mobility is not only linked to the sequence stretch between the WUS homeodomain and the WUS box, but regulation also takes place at the N-and C-terminus. Lacking controls for correct folding of the fusion proteins, we cannot pinpoint changes in mobility, even considering the single, 17 amino acid substitution alleles, to specific sequence stretches. It was however striking that while for the transition from L3 to L2, we saw differences in mobility between WUSΔbox and several substitution alleles, we could not see such differences for the transition from L2 to L1, again suggesting that regulation of WUS mobility is a property of the L3.

In summary, by bringing together massive imaging, careful computational quantification and mechanistic computational modelling, our study reveals that cell-to-cell mobility of the stem cell inducing WUS transcription factor is controlled by a tug of war between uniform active transport and protein retention in the L3. This results in a mobility checkpoint at the L3 to L2 boundary, which coincides the first cell-cell transition outside of the *WUS* RNA expression domain, but is already inside the stem cell domain. Future studies on cell type specific interactors of WUS will be required to uncover the mechanisms underlying this highly specific protein behavior, which represents one of the key control elements of plant stem cell regulation.

## Material and Methods

### Plant material and growth conditions

All plants were of the Col-0 accession. In general, plants were grown at 23°C, 65% relative humidity, under long day conditions (16 hours light, 8 hours darkness) with red and blue light-emitting diodes (LEDs).

Established plant lines (homozygous, single-insertion lines) were not subjected to antibiotic selection. Instead, seeds were sown on soil, were stratified at 4°C in darkness for 2 days and were transferred to long day conditions for subsequent growth.

Selection of transgenic plants using glufosinate-ammonium (BASTA, Bayer CropScience Deutschland GmbH) was performed on soil soaked with 20 mg/l BASTA solution. After sowing, seeds were stratified at 4°C in darkness for 2 days and were transferred to long day conditions. Subsequently, plants were sprayed with 20 mg/l BASTA solution, starting 6-7 days after germination, every 2-3 days for 3-4 times.

Selection of transgenic plants using hygromycin was performed on ½ Murashige-Skoog selection plates (2.15 g/l MS, Duchefa Biochemie; 0.5 g/l MES, Merck KGaA; pH 5.7; 0.7% phytoagar, Duchefa Biochemie; autoclaved; 25 µg/ml hygromycin, Merck KGaA). Plates with surface-sterilized seeds were stratified at 4°C in darkness for 2 days. Afterwards, plates were subjected to a light pulse of 6 hours and were subsequently incubated in darkness for 3 days followed by incubation in light, at long day conditions, for another 3 days.

### Transgenes

All constructs created in this study were assembled using the GreenGate cloning system ^27^. Constructs were transformed ^43^ in Col-0 plants with either wildtype background or *wus-1* mutant background (GK870H12) or in a stable line expressing *pUBQ10:3xmCherry-NLS* in Col-0 wildtype background ^33^. The *wus-1* rescue line GD44 has been described previously ^2^.

### Meristem preparation and confocal microscopy

Meristem preparation and confocal microscopy was performed as has been described previously ^29^ using a Nikon A1 confocal microscope with an Eclipse Ni-E upright stand, a CFI Apochromat long working distance (LWD) 25x 1.1 NA water immersion objective and a 4-channel detector unit including two GaAsP-PMTs, which were used for imaging GFP and mCherry.

### Quantitative image analysis

For the quantification of layer specific distribution of fluorescently tagged WUS protein in the SAM, based on the analysis of fluorescence intensity within a three-dimensional image stack, we developed our ‘Image Transformation and Quantification Tool’ (ITQT) as a plugin to be implemented in the Fiji software package (v1.52p)^44^. In brief, ITQT helps to identify the domain of interest, performs a polar transformation to align cells of the same cell layer, therefore compensating for the curvature of the tissue, and analyzes the total fluorescence intensity within the three-dimensional column defined by the domain of interest. More detailed information on the function of the ITQT plugin can be found in the supplementary methods.

Automated, threshold-based assignment of the center point of the circular analysis domain column was used for all constructs under control of the *WUS* promoter: The ‘Upper Threshold’ was set to an intensity value of 4095; the ‘Lower Threshold’ was preset to 35% of the ‘Upper Threshold’, but was modified as seemed necessary during analysis; the radius of the 3D-median filter was set to 1. Manual assignment of the center point of the circular analysis domain column was used for all constructs under control of the *ML1* promoter: The ‘Periphery Width’ for peripheral analysis was set to 10 µm. All data was analyzed with the ‘Analysis Domain Diameter’ set to 20 µm and the ‘Circle Point Selection Delay’ set to 200 ms.

### Mathematical Modeling

Detailed information on the mechanistic mathematical model for protein distribution in the shoot apical meristem including code can be found in the supplementary methods.

### Data visualization

Micrographs were processed and edited using the Fiji image processing software ^44^. Data plots were generated using the PlotsOfData ^45^ and SuperPlotsOfData ^46^ web applications. Plot style and appearance were adjusted using the open-source software Inkscape or the most recent version of Adobe Illustrator.

## Data Availability

All material described here will be available upon request. Microscopy data, including raw data and data derived from ITQT analysis, that was generated and used in this study will be available at BioImage Archive. The code of the ITQT plugin as well as the code of the mechanistic mathematical model will be available at GitHub.

---

**Figure S1.**
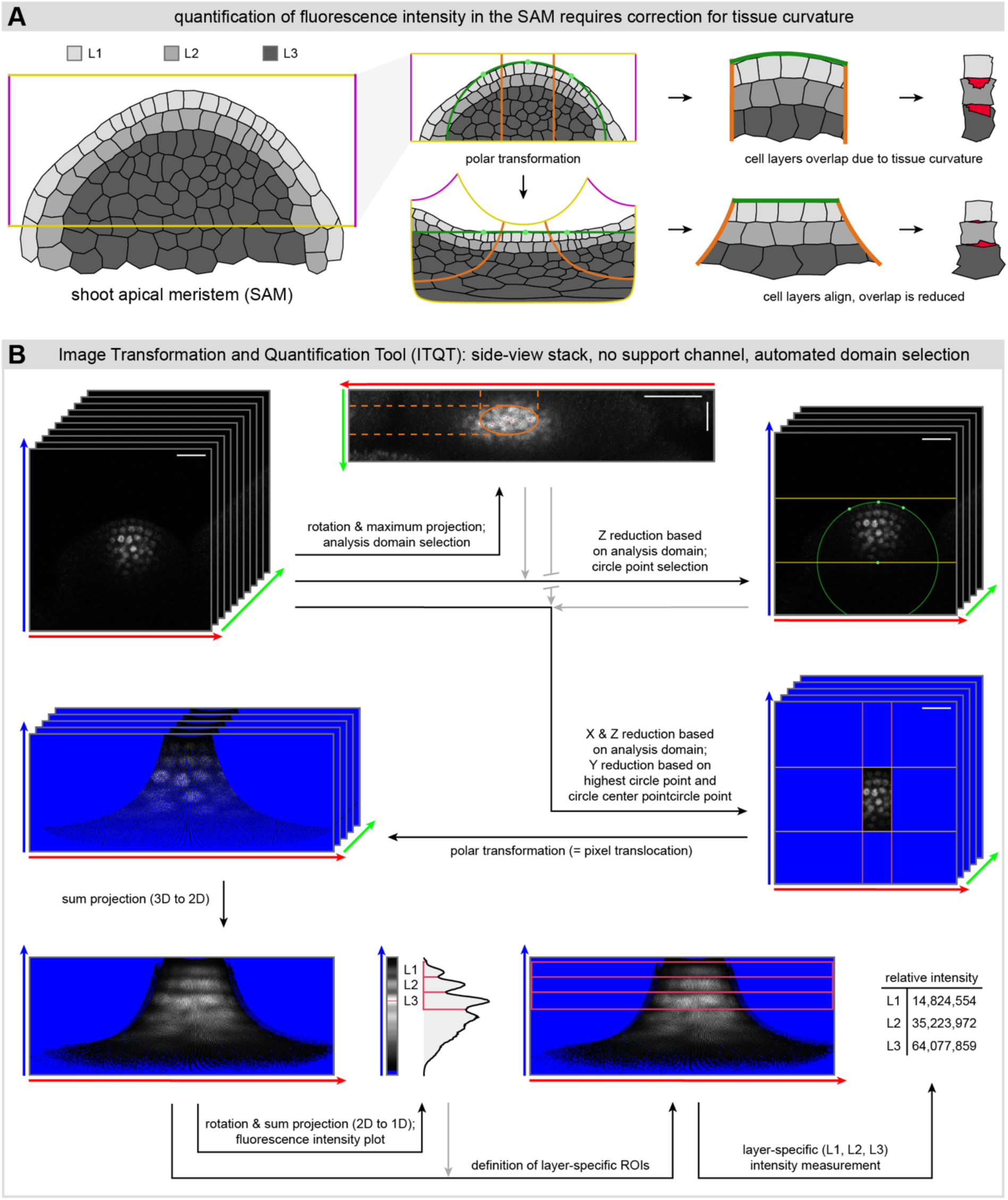
Alignment of meristematic cell layers using ITQT. **(A)** Schematic representation of the importance of cell layer alignment in the shoot apical meristem. **(B)** Schematic workflow of image analysis using the ITQT tool on an exemplary image stack (imaged from a side-view perspective; no secondary support-channel for identification of meristematic cell layers; processed with semi-automated, threshold-based detection of the analysis domain; no additional analysis of a peripheral domain).

**Figure S2.**
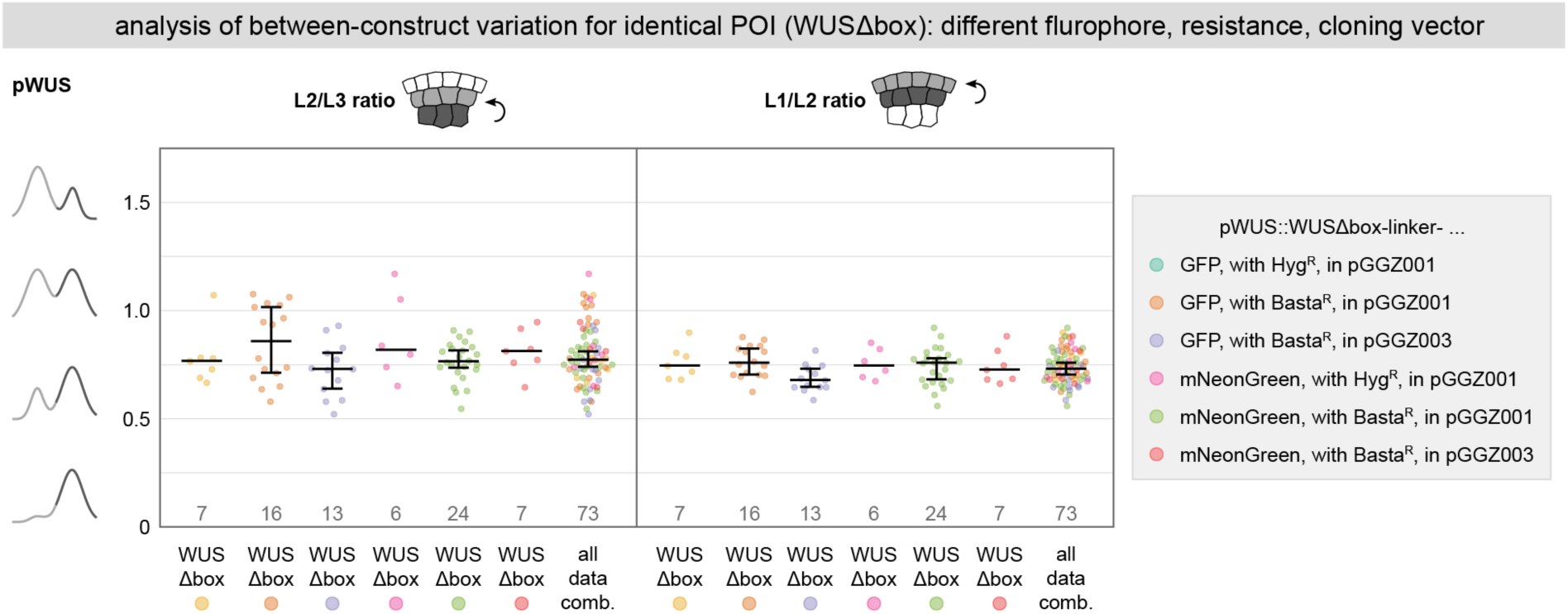
Comparison of between-construct variation of basal to apical mobility for identical POI (WUSΔbox) using different construct designs. Quantitative analysis of upwards mobility of WUSΔbox fusion proteins expressed from the *WUS* promoter (T1 plants) using the ITQT tool. Vectors differ in the fluorophore used for tagging WUSΔbox, the plant resistance gene and the cloning vector (backbone), as is indicated by the different colors. Lines represent population median values with a 95% confidence interval. Sample numbers are indicated below each population.

**Figure S3.**
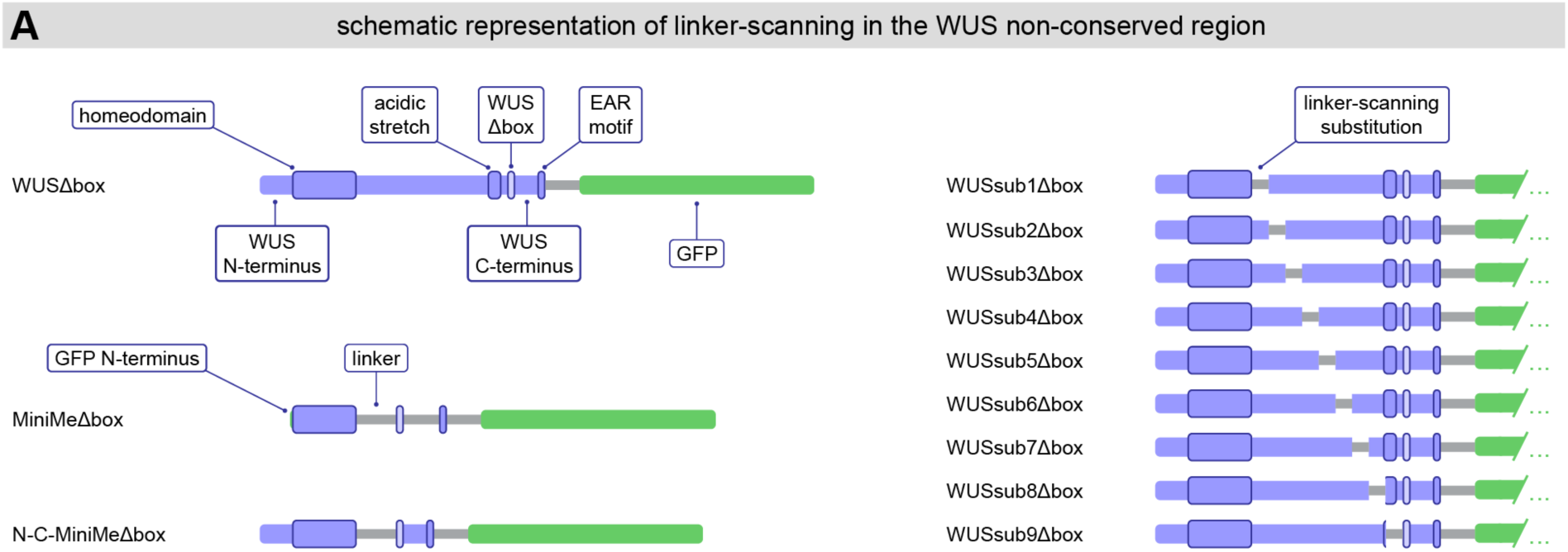
Schematic overview of different linker-substitution WUS fusion proteins. The MiniMeΔbox linker-substitution fusion protein has been previously published ^2^. The N-C-MiniMeΔbox linker-substitution fusion protein reintroduces the WUS N-and C-terminus to the MiniMeΔbox protein while retaining the substitution of a linker for the non-conserved region between the WUS homeodomain and the WUS (Δ)box. WUSsub1Δbox to WUSsub9Δbox fusion proteins were created for linker-scanning of the non-conserved region between the WUS homeodomain and the WUS (Δ)box.

## 1 Model

In this section, we derive the modeling framework. The model considers the three epidermal layers L1, L2 and L3. We use a multi-compartment approach, where the cytoplasm of each layer corresponds to one compartment. The high number number of molecules and the experimental readout with one fluorescence value per layer justify this choice. Molecules of the considered species (WUS, 2xGFP, MiniMi, WOX13, NESWUS) are produced in L1, mediated by the promotor pML1, or in L3, mediated by the WUS promotor pWUS. Molecules in the cytoplasma can be transferred to the nucleus and vice versa. Molecules in the nucleus can bind to and unbind from proteins. Free molecules in the nucleus and protein-bound molecules in the nucleus of each layer are modeled by separate compartments. Free molecules in the cytoplasm can move to the adjacent layers. The term describing the movement accounts for passive diffusion and active transport. The considered processes are summarized in Fig. 1.

### 1.1 L1

In L1 we consider the production of molecules under the promotor *pML*1, degradation of molecules in the cytoplasm, transfer of molecules from the cytoplasm to the nucleus and back. In addition to this molecules in the nucleus can bind to protein and unbind from protein. We consider the following variables:

*u*_1_ : concentration in cytoplasm in L1
*u^n^* : free concentration in nucleus of L1
*u^n,p^* : protein-bound concentration in nucleus of L1

Processes in L1 are quantified by the following parameters:

*m_L_*_2*L*1_ : mobility rate L2 *→* L1
*m_L_*_1*L*2_ : mobility rate L1 *→* L2
*d*_1_ : degradation rate in cytoplasm of L1
*κ* : transport rate from cytoplasm to nucleus *λ* : transport rate from nucleus to cytoplasm *α*_1_ : protein binding rate in nucleus of L1
*β*_1_ : protein unbinding rate in nucleus of L1
*p*_1_ : production mediated by pML1

Volume differences of nucleus and cytoplasm are incorporated in the transport rates.

The concentration of free molecules in the cytoplasm *u*_1_ is described by the following ODE:

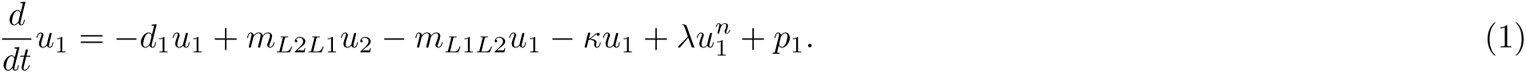

The concentration of free molecules in the nucleus *u^n^*is described by the following ODE:

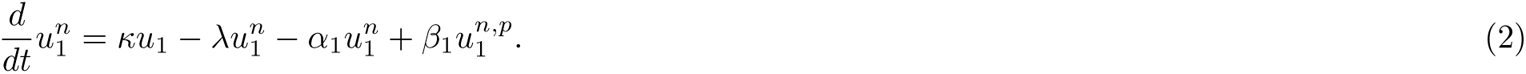

The concentration of protein-bound molecules in the nucleus *u^n,p^* is described by the following ODE:

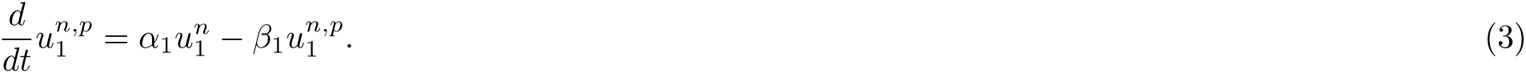

**Figure 1:**
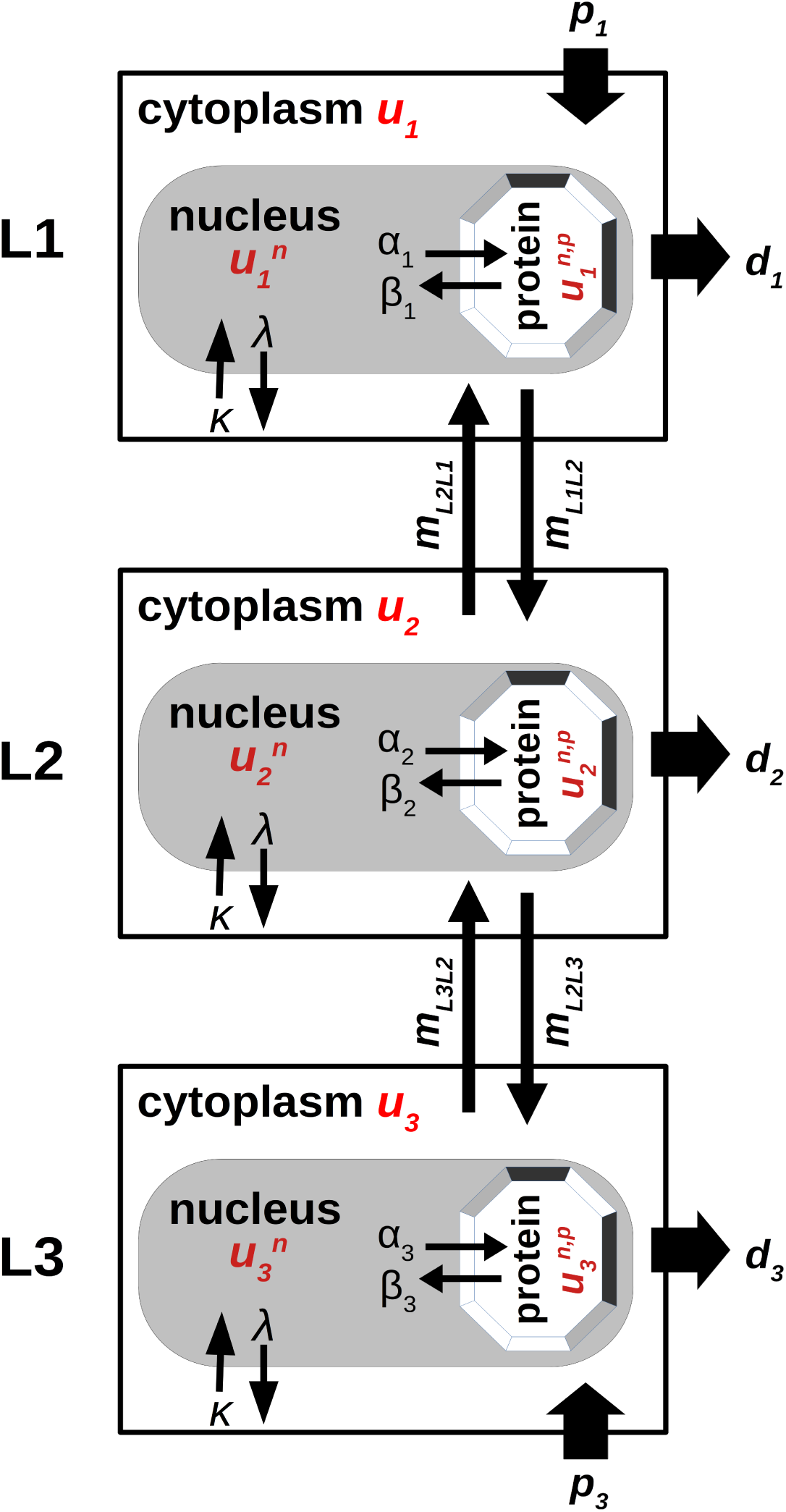
Overview of the model. Process parameters are indicated in black, model variables in red.

### 1.2 L2

In L2 we consider degradation of molecules in the cytoplasm, transfer of molecules from the cytoplasm to the nucleus and back. In addition to this, molecules in the nucleus can bind to protein and unbind from protein. We consider the following variables:

*u*_2_ : concentration in cytoplasm in L2
*u^n^* : concentration in nucleus in L2
*u^n,p^* : protein-bound concentration in nucleus of L2

In addition to the parameters introduced above, we use the following parameters to quantify the processes in L2:

*m_L_*_3*L*2_ : mobility rate L3 *→* L2
*m_L_*_2*L*3_ : mobility rate L2 *→* L3
*d*_2_ : degradation rate in cytoplasm of L2
*α*_2_ : protein binding rate in nucleus of L2
*β*_2_ : protein unbinding rate in nucleus of L2

The concentration of free molecules in the cytoplasm *u*_2_ is described by the following ODE:

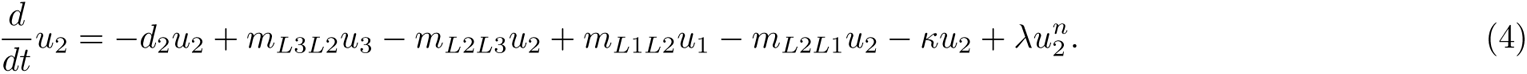

The concentration of free molecules in the nucleus *u^n^*is described by the following ODE:

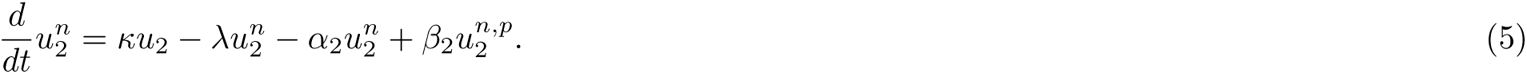

The concentration of protein-bound molecules in the nucleus *u^n,p^* is described by the following ODE:

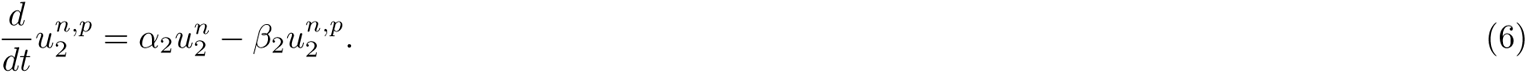

### 1.3 L3

In L3 we consider the production of molecules under the promotor *pWUS*, degradation of molecules in the cytoplasm, transfer of molecules from the cytoplasm to the nucleus and back. In addition to this, molecules in the nucleus can bind to protein and unbind from protein. We consider the following variables:

*u*_3_ : concentration in cytoplasm in L3
*u^n^* : free concentration in nucleus of L3
*u^n,p^* : protein-bound concentration in nucleus of L2

In addition to the parameters introduced above, we use the following parameters to quantify the processes in L3:

*d*_3_ : degradation rate in L3
*α*_3_ : protein binding rate in nucleus of L2
*β*_3_ : protein unbinding rate in nucleus of L2
*p*_3_ : production mediated by pWUS

The concentration of free molecules in the cytoplasm *u*_3_ is described by the following ODE:

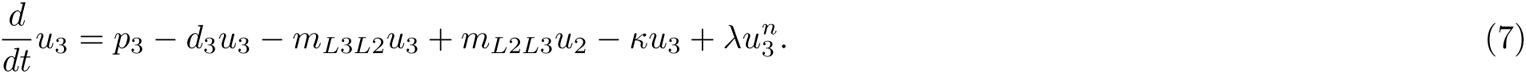

The concentration of free molecules in the nucleus *u^n^*is described by the following ODE:

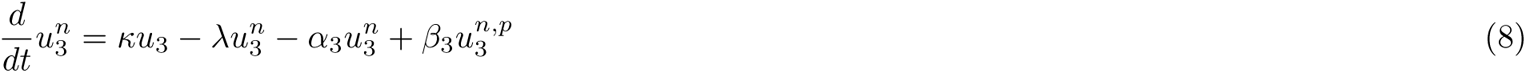

The concentration of protein-bound molecules in the nucleus *u^n,p^* is described by the following ODE:

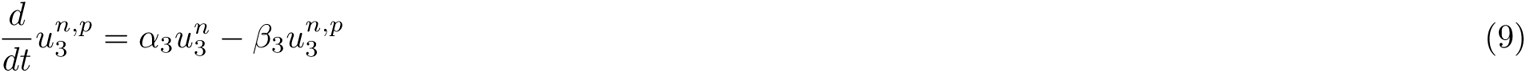

## 2 Equilibrium state

We assume that the system is in an equilibrium state. We explicitly calculate the equilibrium state.

We denote the equilibrium concentrations by a bar. In the equilibrium state it holds due to equations (2), (3), (5), (6), (8) and (9):

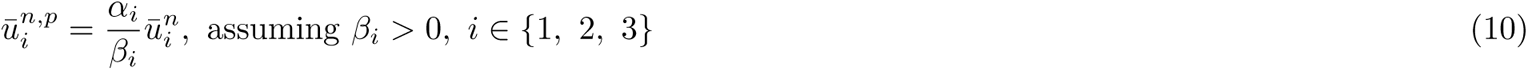

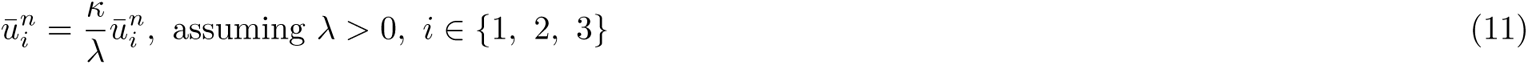

We observe that the equilibrium concentrations depend only on the ratios 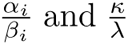 but not on the individual parameters. For this reason we fix *β_i_* = 1 and *λ* = 1 for the parameter fitting.

Furthermore it holds due to equation (1):

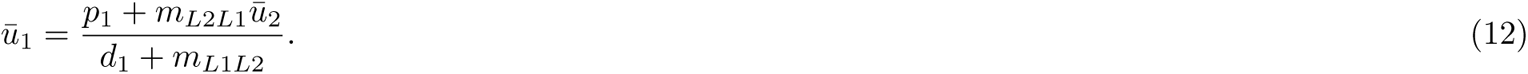

[]Equations (4) and (12) imply

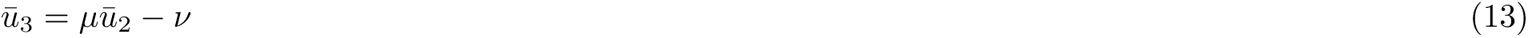

with

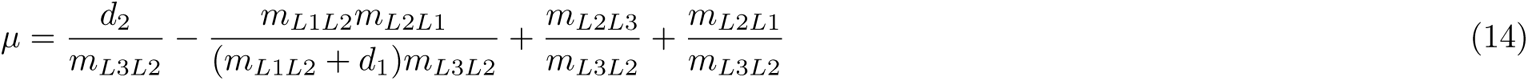

and

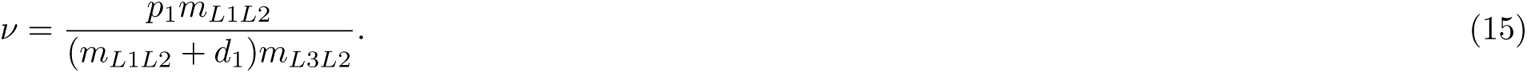

Equations (1) and (13) imply

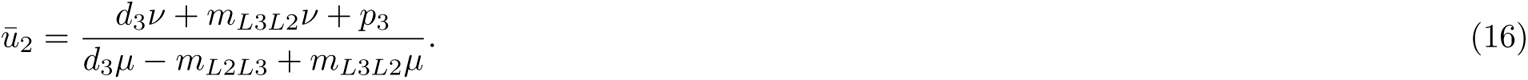

We calculate the steady state using equations (16), (13), (12), (11) and (10) and implement it accordingly in the matlab scripts.

## 3 Parameter fitting

We use the MATLAB function fmincon to estimate parameters. We assume that the system is in equilibrium. We consider a weighted least square cost functional, where the inverse of the empirical variances are used as weights. We compare 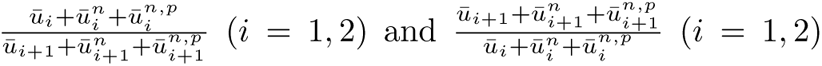 to the measured fluorescence ratios. For each experiment we use 200 multi-starts with random initial guesses obtained from latin hypercube sampling. We fix the expression rates *p*_1_ and *p*_3_ at 1. Since the nucelus-cytoplasma-ratio 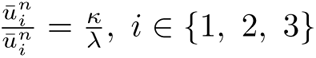 depends only on the ratio of *κ* and *λ*, we set *λ* = 1. Analogously, we set *β_i_* = 1*, i ∈ {*1, 2, 3*}*. The parameters in the optima are unique up to a scaling constant.

## Notes

### Competing Interest Statement

The authors have declared no competing interest.

